# Invariant Natural Killer T cells control positively and negatively the development of hepatocellular carcinoma

**DOI:** 10.1101/2024.08.29.610221

**Authors:** Maria Papanastasatou, Marianthi Gioulbasani, Evangelia Nakou, Alexandros Galaras, Teresa Rubio-Tomás, Iannis Talianidis, Aristides Eliopoulos, Pantelis Hatzis, Mihalis Verykokakis

**Affiliations:** Institute for Fundamental Biomedical Research, BSRC Alexander Fleming, Vari, Greece; Department of Biology, Medical School, National and Kapodistrian University of Athens, Athens, Greece; Department of Biochemistry and Biotechnology, University of Thessaly, Larissa, Greece; Lineberger Comprehensive Cancer Center, University of North Carolina, Chapel Hill, NC, USA; Intelligencia AI, Athens, Greece; Institute of Molecular Biology and Biotechnology, Foundation for Research and Technology-Hellas, Heraklion, Greece

## Abstract

The liver routinely encounters antigens from the gut, triggering pro-inflammatory responses. Unresolved inflammation can lead to liver damage, steatosis, fibrosis, cirrhosis, and eventually hepatocellular carcinoma (HCC). HCC is influenced by various immune cells, including invariant natural killer T (iNKT) cells, which exhibit both innate and adaptive immunity traits. Here, we examined iNKT cell dynamics in a diethyl-nitrosamine (DEN)-induced HCC mouse model. We observed a significant reduction in iNKT cell numbers in HCC livers due to apoptosis and impaired cytokine production. *CD1d*-deficient mice, which lack iNKT cells, displayed delayed tumor initiation and lower tumor and foci number. However, these tumors were larger in size and characterized by enhanced proliferation and immunosuppression. Interestingly, adoptive transfer of healthy iNKT cells post-tumor establishment reduced tumor burden, highlighting their potential therapeutic role. Our findings suggest that iNKT cells contribute to early HCC development, while in later stages they help to control tumor growth, thus underscoring their complex role in liver carcinogenesis. Further understanding of iNKT cell functions may inform novel immunotherapeutic strategies for HCC management.

## Introduction

Conventionally, the liver is acknowledged as the primary metabolic hub in the body, carrying out a multitude of functions such as carbohydrate metabolism, nutrient absorption and storage, biosynthesis of biochemical compounds, lipid metabolism, and detoxification^1^. In healthy individuals, the liver regularly encounters various foreign microbial and dietary antigens originating from the gut. These antigens can be detected by receptors present on hepatic cells, triggering a pro-inflammatory response. Nevertheless, an uncontrolled inflammatory reaction during homeostasis could prove detrimental, ultimately culminating in liver injury and inflammation^2^. The inability to effectively resolve inflammation is associated with the development of liver damage, marked by steatosis and fibrosis. This condition can advance to cirrhosis and eventually lead to hepatocellular carcinoma (HCC)^3^.

HCC constitutes approximately 80% of primary liver cancer and is a significant cause of mortality in Western countries^4^. The development of HCC is linked to chronic inflammation resulting from viral infections or other environmental factors. Despite recent progress, existing therapies for HCC yield limited responses in patients. Hence, novel strategies are required to address the current therapeutic needs.

HCC is associated with the functions of a diverse array of immune cells, including invariant natural killer T cells (iNKT), cytotoxic CD8 T cells, CD4 T helper (Th) cells, regulatory T cells (T_reg_), myeloid-derived suppressor cells (MDSC), NK cells (NK), and dendritic cells (DC)^5^. Notably, in HCC, dendritic cells exhibit a diminished capacity to recognize and present cancer antigens to T cells^6^. Furthermore, MDSCs suppress the function of immune cells^7^ and foster the expansion of T_regs_^8^. T_regs_ negatively impact T cell proliferation by suppressing IFNγ secretion by T cells^9^ and the responses of NK cells^10^. Moreover, CD8 T cells demonstrate dysfunction in HCC, concomitant with a significant reduction in the number of CD4 T cells^10^.

Invariant NKT cells, which constitute a rare subset of T cells in both humans and mice, exhibit characteristics of both innate and adaptive immunity. iNKT cells, similar to conventional T cells, undergo development within the thymus from T cell progenitors and generate a functional oligoclonal T cell receptor (TCR)^11,12^. iNKT cells originate from double positive (DP) precursors expressing CD4 and CD8^11^. They bear the characteristic iNKT TCRα (Vα14- Jα18 in mice, Vα24-Jα18 in humans), following random genomic rearrangement of the *Tcrα* locus^11^ , which pairs with a limited number of Vβ chains^11^. Within the thymus, iNKT cells acquire their innate-like properties, without foreign antigen stimulation, and differentiate in at least three mature subsets, distinguished by the constitutive expression of specific transcription factors associated with conventional Th lineages: iNKT1 cells (predominant in the liver) express the Th1-related transcription factor TBET, iNKT2 cells express GATA3, and iNKT17 cells express RORγt^13^. Predominantly found in the liver (comprising 10-15% of total lymphocytes), iNKT cells recognize lipid antigens - rather than peptides - presented by CD1d. CD1d is expressed in various cells, including antigen-presenting cells and hepatocytes^14^. Upon activation, iNKT cells rapidly produce large amounts of diverse cytokines and chemokines orchestrating the early phase of the immune response^15,16^.

In cancer immunotherapy strategies, iNKT cells offer advantages over conventional T cells. Both pre-clinical and clinical evidence supports the protective role of donor iNKT cells against graft-versus-host disease (GVHD)^17^. iNKT cells possess the ability to directly eliminate target cells expressing CD1d, including cancer cells, utilizing perforin^18^ and TNFα-mediated cytotoxic pathways^19^. Additionally, iNKT cells rapidly produce cytokines and chemokines, activating both innate and adaptive immune cells, thereby influencing the tumor microenvironment^20^. Finally, iNKT cells kill tumor-associated macrophages (TAMs)^21^, which promote tumor progression, and alleviate immune suppression mediated by CD1d^+^ MDSCs^22^.

iNKT cells serve as targets in immunotherapeutic approaches for various cancer types. In a melanoma model, activated iNKT cells demonstrated a potent inhibitory effect on tumor metastasis by exerting strong cytolytic activity against metastasized tumor cells in the liver^23^. Furthermore, intravenous injection of IL-12 in erythroleukemia resulted in suppressed tumor growth and metastasis, attributed to the direct cytotoxicity mediated by iNKT cells^24^. Similarly, intravenous administration of alpha-GalactosylCeramide (α-GalCer), which strongly activates iNKT cells, in a colon hepatic metastasis adenocarcinoma mouse model inhibited tumor growth in the liver through the activation of NK and T cells^25^. Notably, iNKT cells expressing chimeric antigen receptors (CARs) demonstrated significant antitumor activity, sustained tumor regression, and improved survival rates^26,27^.

While iNKT cells have been found to promote liver damage^28,29^, their specific involvement in HCC mechanisms remains elusive^30,31^. In orthotopic mouse hepatomas, iNKT cell activation with α-GalCer administration suppressed tumor growth through NK cell activation^32^, suggesting a strong anti-tumor iNKT cell function. Consistent with that, adoptive transfer of iNKT cells that were activated with HCC-derived antigens suppressed HEP3B-mediated liver tumorigenesis^33^. However, these liver cancer mouse models lack the component of chronic inflammation and fibrosis that commonly precedes HCC development. Indeed, adoptive transfer of iNKT cells (but not conventional CD4

T cells) enhanced fibrogenesis, which is associated with HCC^31,34^, indicating that, overall, iNKT cells may also contribute to HCC promotion. iNKT cell depletion through administration of anti-NK1.1 resulted in aggravated liver tumorigenesis in a β-catenin-mediated model of hepatic inflammation^35^, thus indicating a potential anti-inflammatory role for iNKT cells; however, anti-NK1.1 treatment also depletes NK cells, which have strong anti-tumor activity. These findings underscore possible multiple roles for iNKT cells during the course of HCC pathogenesis, suggesting context- and possibly time-dependent effects that warrant further investigation.

Here, we induced HCC with characteristics of chronic inflammation, similar to human HCC, with diethyl-nitrosamine (DEN) treatment in mice^36^. Therefore, this HCC mouse model enables monitoring of the functions of iNKT cells during the full course of HCC development, in a comprehensive manner. We found that the iNKT cell number decreased in HCC, due to extensive apoptosis, indicating a potential dysregulation of their population dynamics within the tumor microenvironment. Furthermore, cytokine production by the remaining iNKT cells was impaired in HCC, thus compromising their ability to mount potent anti-tumor responses. Mice that were genetically modified to lack iNKT cells were characterized by reduction in tumor and foci numbers, while the tumor volume increased compared to *WT* mice, implying a nuanced role for iNKT cells in HCC development. Moreover, the absence of iNKT cells ameliorated liver damage during HCC initiation, suggesting their involvement in the early stages of disease pathogenesis. However, iNKT cell loss also led to alterations in the tumor microenvironment, including increased T_reg_ cell number and enhanced immunosuppression, ultimately promoting hepatocyte cell proliferation. Interestingly, adoptive transfer of naïve healthy iNKT cells after tumor establishment (6 months post-DEN injection) led to a decrease in the number of foci and tumors, along with a reduction in tumor volume in *WT DEN* mice, while adoptive transfer of iNKT cells in earlier stages (1 and 3 months after DEN administration) slightly enhanced the development of HCC. Therefore, adoptive transfer of iNKT cells emerges as a potential therapeutic strategy to suppress liver carcinogenesis after tumor establishment, highlighting the importance of understanding the functions of iNKT cells in HCC management.

## Results

### The number of iNKT cells decreased in HCC

To begin to understand the function of iNKT cells in HCC, we induced liver tumors with the administration of 25 mg/kg diethyl-nitrosamine (DEN) in 15-day old male *C57BL/6 WT* mice^36^. The DEN-induced mouse HCC is highly immunogenic and develops in the background of chronic inflammation; therefore, it is best suited to study anti-tumor immune responses and the contributions of immune cell populations, such as iNKT cells^37^. Mice treated with DEN developed multiple macroscopically visible tumors (>3 mm in size) and smaller foci (<3 mm) in the liver 38-42 weeks post-treatment, while histological analysis showed extensive liver damage and focal hyperplasia, and extensive recruitment of lymphocytes around cancer lesions (Figure 1A). In addition, we noticed an increase in liver-to-body weight ratio compared to untreated *WT* mice, consistent with the presence of enlarged livers and HCC development (Figure 1B). Real-time PCR revealed an increase in the expression of the HCC marker genes *Afp* and *Gpc3* in *WT DEN* compared to *WT CTRL* mice (Figure 1C). Consistent with the histological findings, flow cytometry analysis unveiled an increased number of lymphocytes in mice with HCC compared to healthy mice (Figure 1D).

**Figure 1.**
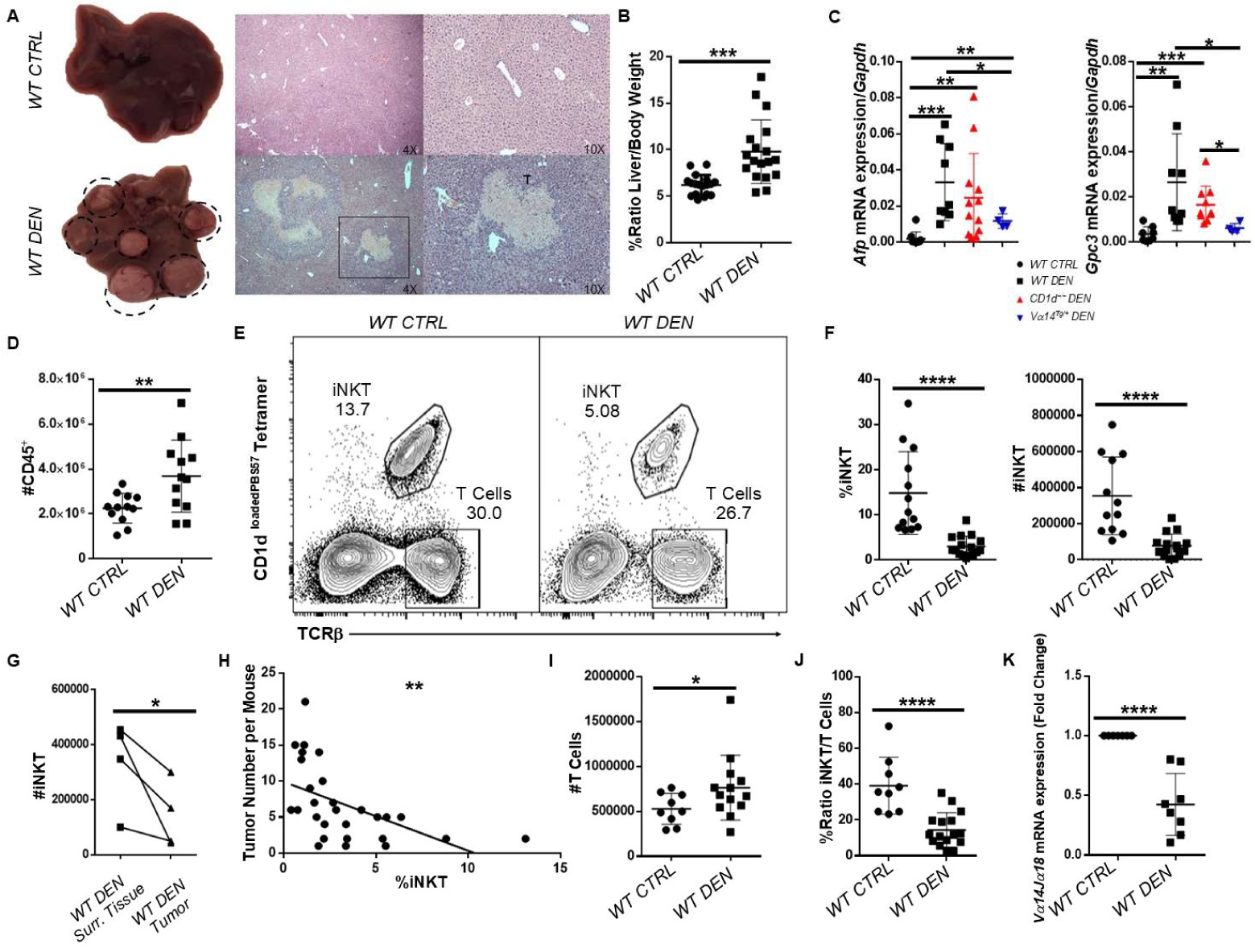
The number of iNKT cells decreased in HCC. **A**. Liver images from WT CTRL (top) and WT DEN (bottom) mice. H&E histology 38-42 weeks after DEN treatment in 4X and 10X magnification. **T**, tumor. **B** Increased %Ratio Liver/Body weight in WT DEN mice. **C** Scatter plots show the mRNA expression of Afp and Gpc3 in the liver from WT CTRL, WT DEN, CD1d^−/−^ DEN, and Vα14^Tg/+^ DEN mice. **D** Number of CD45^+^ cells are shown. **E** FACS plots of total liver lymphocytes from WT and WT DEN mice showing the percentage of iNKT (Tetr^+^TCRβ^+^) and conventional T (Tetr^−^TCRβ^+^) cells. **F** Scatter plots showing the percentage and number of iNKT cells. **G** Scatter plot shows the number of iNKT cells in surrounding tissue and in tumor from WT DEN mice. **H** Scatter plot shows the correlation of the frequency of iNKT cells (X axis) with the number of tumors per mouse (Y axis) in WT DEN mice. p< 0.01, R^2^= 0.2915. **I** Scatter plot shows the number of T cells. **J** Scatter plot shows the %Ratio iNKT/T cells in the liver. **K** Scatter plot shows the Vα14Jα18 mRNA expression in WT and WT DEN mice. Bars represent mean ± SD from 28 independent experiments for macroscopic analysis, 12 independent experiments for mRNA of Afp and Gpc3 and flow cytometry analysis, 4 independent experiments for flow cytometry analysis between surrounding tissue and tumor and 7 independent experiments for mRNA of Vα14Jα18 transcript with one WT and one or two WT DEN mice each. *, p<0.05, **, p<0.01, ***, p<0.001, ****, p<0.0001 by t test.

We next to sought to examine the presence of iNKT cells in HCC livers, using CD1d tetramers loaded with the alpha-GalactosylCeramide analogue PBS57. Our flow cytometry analysis revealed a significant decrease in both the percentage and number of iNKT cells (Tetr^+^TCRβ^+^) in *WT DEN* mice compared to *WT CTRL* mice in the liver (Figure 1E, F). Importantly, the number of intratumoral iNKT cells was also reduced compared to surrounding healthy tissue from mice with HCC (Figure 1G). Additionally, the frequency of iNKT cells in the tumor was anti-correlated with the number of developed tumors in *WT DEN*-treated mice (Figure 1H), indicating a potential beneficial role of iNKT in tumor control. No observable changes were detected in the frequency and number of iNKT cells in the spleen (Suppl. Fig. 1A, B), indicating a specific loss of hepatic iNKT cells during HCC development in mice. In contrast, we found an increase in the number of conventional T cells in the livers of mice with HCC compared to healthy mice (Figure 1I); consequently, the iNKT/T cell ratio decreased in *WT DEN* mice (Figure 1J). Furthermore, the frequency and number of macrophages (CD11b^+^F4/80^+^) increased in *WT DEN* compared to *WT CTRL* mice (Suppl. Fig. 1C, D).

Because iNKT cell activation results in downregulation of surface TCR expression, flow cytometry may underestimate the presence of iNKT cells, in case they are activated in HCC. To validate the reduction of iNKT cells in HCC, we assessed the expression levels of the *Vα14Jα18* transcripts, which is unique to iNKT cells. As shown by qPCR, *Vα14Jα18* mRNA levels in the liver were significantly lower in *WT DEN* compared to *WT CTRL* mice (Figure 1K). Collectively, our analysis showed that liver iNKT cells were less abundant in mice with HCC, although the total number of conventional T cells and macrophages increased.

### The functionality of iNKT cells was impaired in HCC

We next investigated whether iNKT cells from the HCC microenvironment are competent to mount an effective immune response. To address this, we examined the expression of CD69, which is induced upon TCR engagement and is a common activation marker of T cells, on iNKT cells from *WT DEN*-treated mice. Typically, the vast majority of hepatic iNKT cells from healthy mice consistently express CD69, consistent with their pre-activation status (Figure 2A, left panel). In contrast, only approximately 60% of hepatic iNKT cells expressed CD69 in HCC mice (Figure 2A, B), indicating diminished iNKT cell activation. Furthermore, an increased percentage of hepatic iNKT cells expressed high levels of PD1, which is a marker of T cell exhaustion, suggesting that iNKT cells may be dysfunctional in the context of HCC (Figure 2C, D).

**Figure 2.**
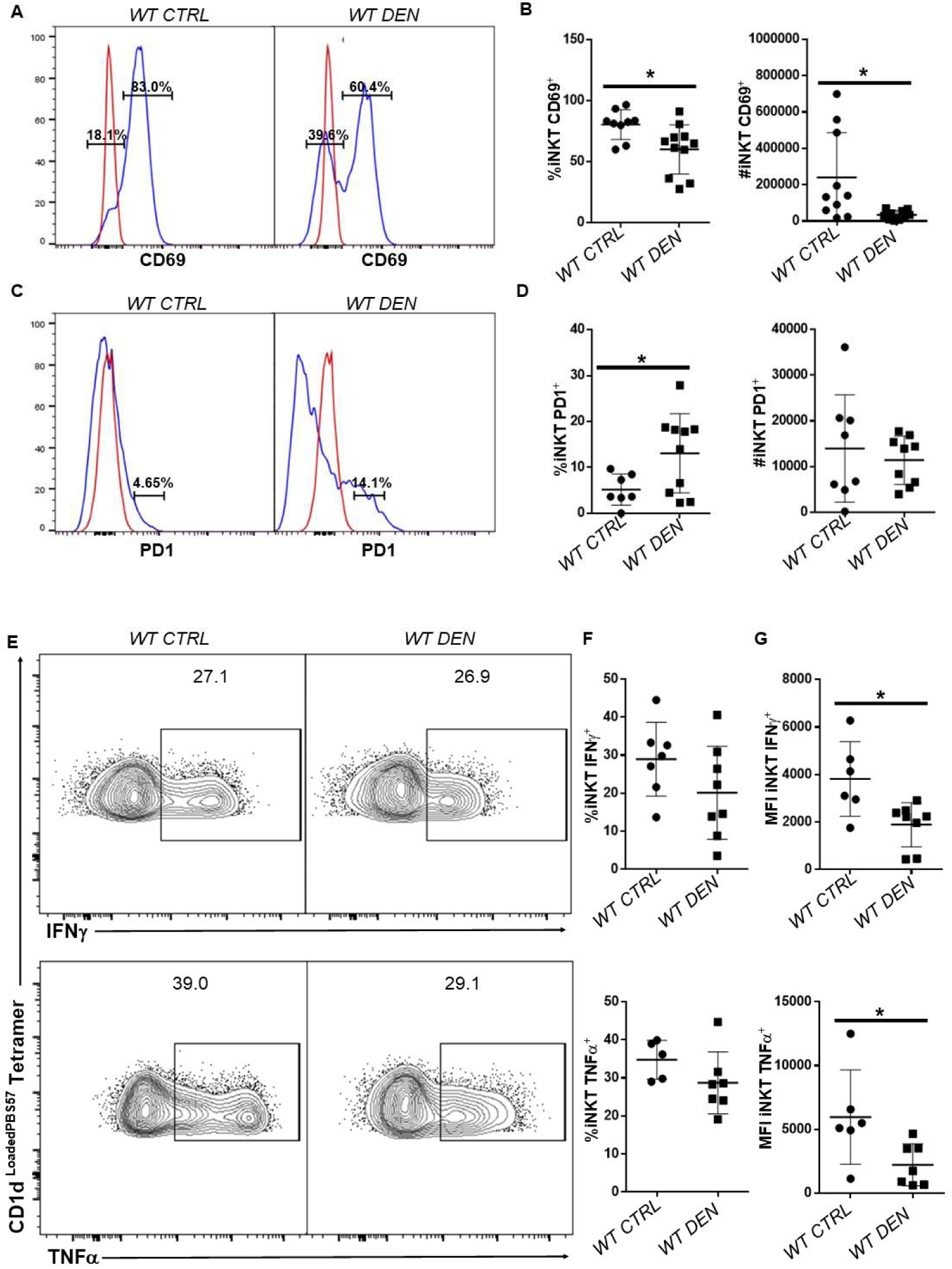
The functionality of iNKT cells was impaired in HCC mice. **A** Histograms show the CD69 expression in iNKT cells from WT and WT DEN mice. Red line shows the FMO of CD69. **B** Scatter plots show the decreased percentage and number of iNKT cells expressing CD69 in HCC. **C** Histograms show the expression of PD1 in iNKT cells from WT and WT DEN mice. Red line shows the FMO of PD1. **D** Scatter plots show PD1 expression in iNKT cells from the indicated mice**. E** FACS plots show the expression of IFNγ and TNFα in hepatic iNKT cells after PMA/ION stimulation in vitro. **F** Scatter plots show the percentage of IFNγ and TNFα in iNKT cells from WT and WT DEN mice. **G** Scatter plots show the MFI of IFNγ and TNFα in iNKT cells from WT and WT DEN mice. Bars represent mean ± SD from 10 independent experiments for activation and exhaustion analysis and 7 independent experiments for the ability of iNKT cells to produce IFNγ and TNFα, with one WT and one or two WT DEN mice each. *, p<0.05 by t test.

To directly examine the ability of iNKT cells to produce cytokines, we stimulated lymphocytes with phorbol 12-myristate 13-acetate (PMA) and ionomycin (ION) *in vitro*. Flow cytometry analysis revealed that the percentage of iNKT cells that were able to produce IFNγ and TNFα was comparable between healthy and HCC mice (Figure 2E and F). Notably however, iNKT cells from HCC livers produced significantly lower amount of IFNγ and TNFα, on a per-cell basis (Figure 2G). Taken together, in addition to the reduced presence of iNKT cells in HCC livers, our results demonstrated that the remaining iNKT cells exhibited impaired activation, increased exhaustion, and overall compromised cytokine production.

### iNKT cells were apoptotic in HCC

To investigate how the gene expression program of iNKT cells is influenced in the context of HCC, we performed global transcriptomic analysis of iNKT cells FACS-sorted from *WT CTRL* and *WT DEN* livers. Our analysis revealed that 156 genes were differentially expressed between iNKT cells from healthy mice and those from HCC mice, with at least a 2-fold change (DESeq, p<0.05) (Figure 3A). Of those, 45 genes were downregulated and 111 were upregulated in iNKT cells from HCC livers. Strikingly, genes upregulated in iNKT cells from HCC mice were associated with cell death pathways and impaired biosynthesis, indicating that iNKT cell fitness in HCC was compromised (Figure 3B). Consistent with this finding, flow cytometry analysis demonstrated a significant increase in the expression of active caspase 3 in iNKT cells from HCC compared to those from healthy mice (Figure 3C and D), which indicates that these cells were apoptotic. In contrast, expression of active caspase 1 in iNKT cells, which is involved in pyroptosis, remained unaltered between healthy and HCC mice (Suppl. Fig. 2). Thus, our observations indicate that the HCC microenvironment may trigger induction of apoptotic pathways in iNKT cells, thus leading to a severe loss of this immune cell subset.

**Figure 3.**
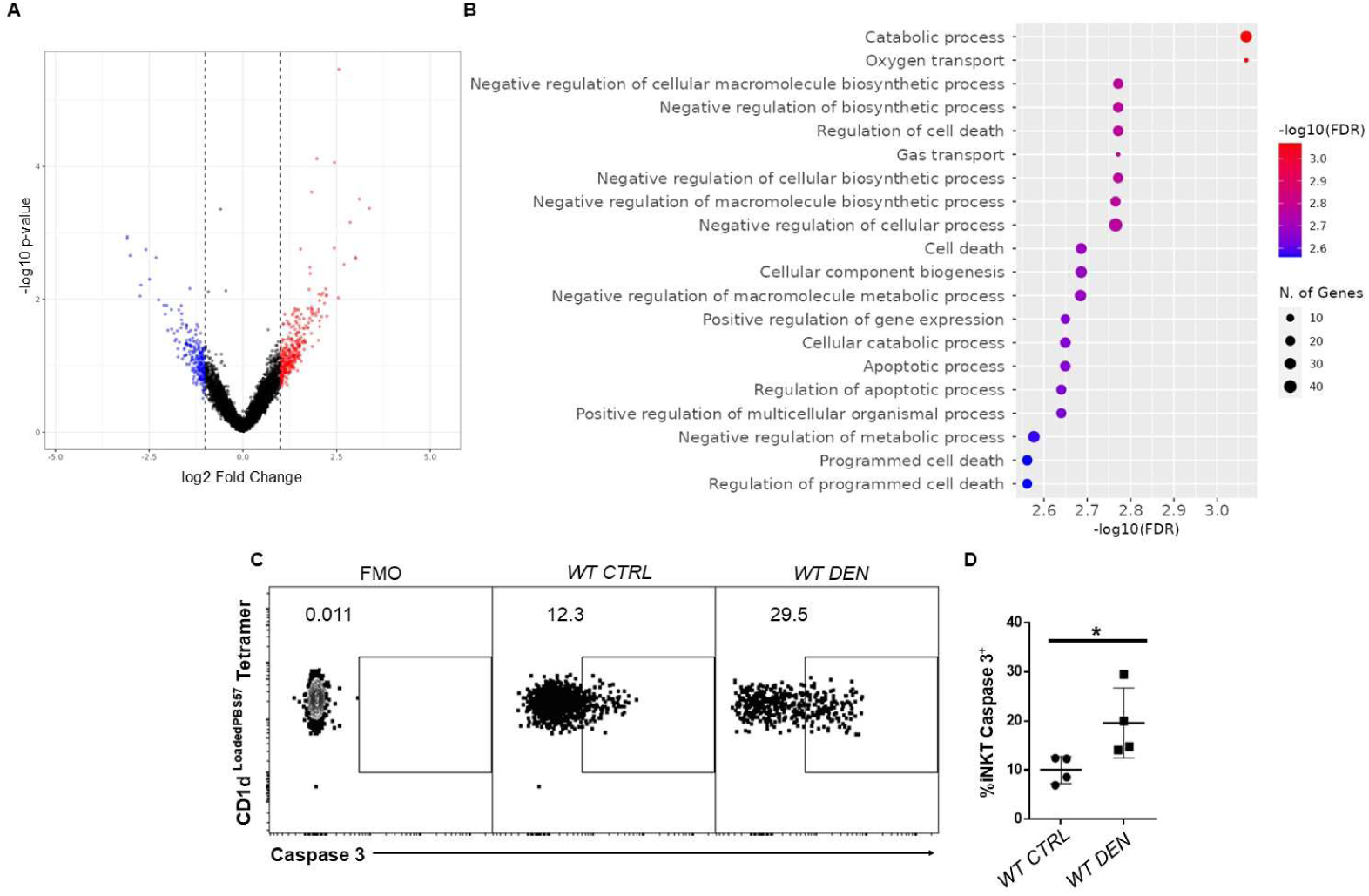
iNKT cells were apoptotic in HCC. **A** Volcano plot showing the upregulated and downregulated genes in iNKT cells from WT CTRL and WT DEN mice. **B** Gene ontology analysis showing the pathways enriched among the genes that were differentially expressed. **C** FACS plots show the expression of active caspase 3 on iNKT cells in WT and WT DEN mice. **D** Scatter plot show the increased expression of active caspase 3 on iNKT cells in WT DEN mice. Bars represent mean ± SD from 3 independent experiments, with one or two WT and one or two WT DEN mouse each. *, p<0.05 by t test.

### *CD1d*-deficient mice showed reduced tumor burden

To investigate the functional requirements for iNKT cells in HCC development, we analyzed *CD1d^−/−^* mice, which lack iNKT and type II NKT cells, due to inhibition of NKT cell positive selection in the thymus^38^. Intriguingly, mice lacking iNKT cells displayed significantly fewer foci and tumors compared to *WT* mice after DEN treatment (Figure 4A, B), although tumor incidence was similar between *WT* and *CD1d^−/−^* mice (Figure 4C). Surprisingly, the few tumors that developed in the absence of iNKT cells had significantly larger volume than tumors developed in *WT* mice (Figure 4D). No significant differences were observed in body and liver weight (Figure 4E and F), and *Afp* and *Gpc3* mRNA levels between *WT* and *CD1d^−/−^* mice after DEN treatment (Figure 1C).

**Figure 4.**
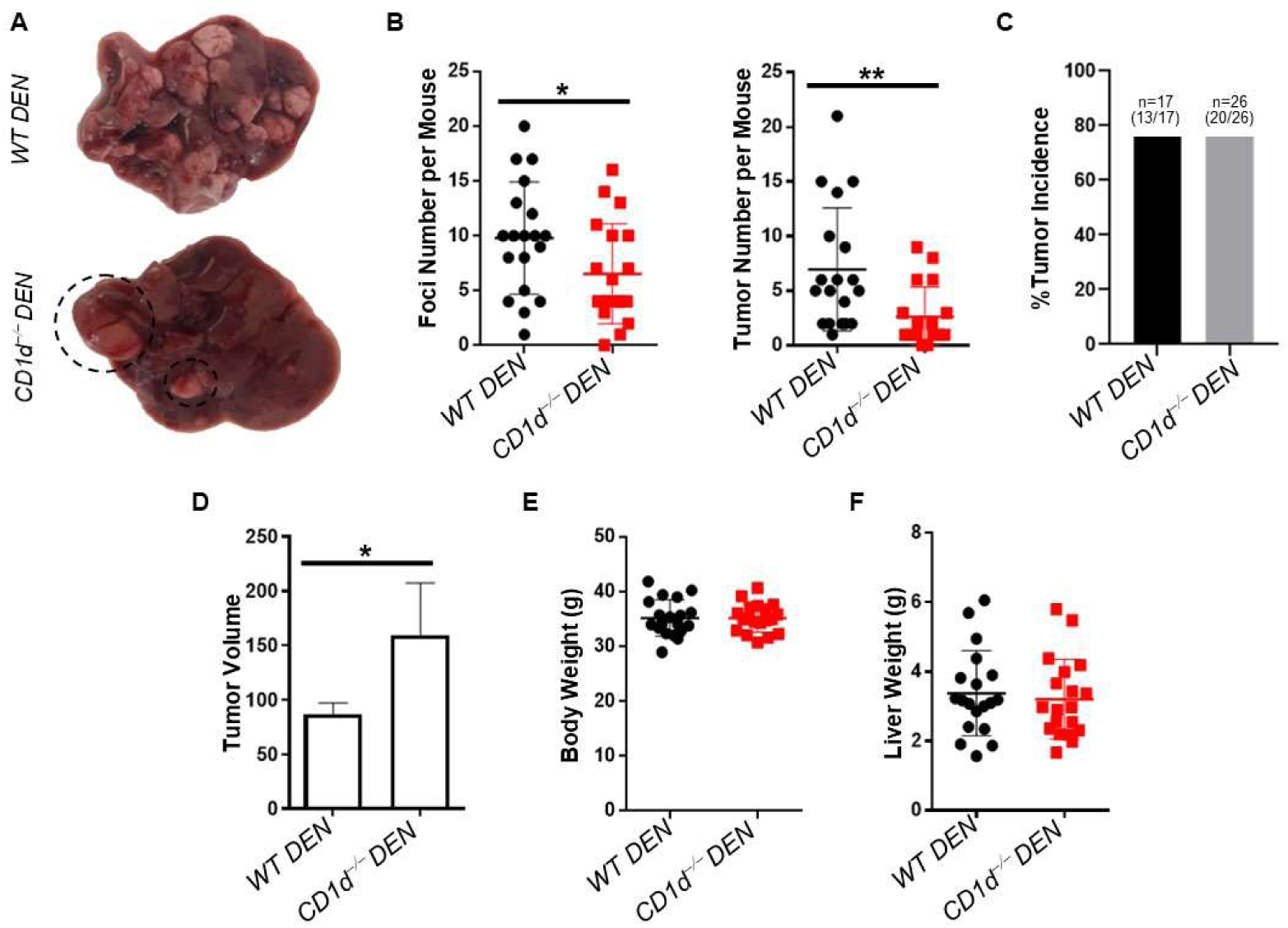
CD1d-deficient mice showed reduced tumor burden. **A** Liver pictures from WT DEN (top) and CD1d^−/−^ DEN (bottom) mice. **B** Scatter plots show foci and tumor number. **C** Bar plot shows the tumor incidence in WT DEN and CD1d^−/−^ DEN mice. **D** Bar plot shows the tumor volume between WT DEN and CD1d^−/−^ DEN mice. **E** Scatter plot shows body weight in WT DEN and CD1d^−/−^ DEN mice. **F** Scatter plot shows liver weight in WT DEN and CD1d^−/−^ DEN mice. Bars represent mean ± SD from 17 independent experiments, with one WT and one or two CD1d^−/−^ DEN mice each 38-42 weeks after DEN administration. *, p<0.05, **, p<0.01 by t test.

Because liver tumors lacked iNKT cells, we hypothesized that excess number of iNKT cells may protect from HCC. To test this hypothesis, we analyzed mice that express a pre-rearranged *Vα14Jα18* TCRα chain (*Vα14^Tg/+^* mice), thus they overproduce iNKT cells from birth^39^. Our analysis revealed that *Vα14^Tg/+^* mice developed the same number of tumors and foci as *WT* mice (Suppl. Fig. 3A and B), although these tumors were smaller in size (Suppl. Fig. 3B). Taken together, these results indicate that iNKT cells contribute to HCC development, although they may restrict the growth of individual tumors.

Consistent with these findings, we found that tumors developed in the absence of iNKT cells showed enhanced proliferation compared to *WT* tumors, based on Ki67 immunofluorescence analysis (Suppl. Fig. 4A, B). Additionally, we measured the mRNA expression of *c-Jun*, which has been associated with increased cell proliferation. Our results unveiled that the mRNA levels of *c-Jun* were significantly increased in *CD1d^−/−^ DEN* mice compared to *WT DEN* mice (Suppl. Fig. 4B).

Moreover, we examined the mRNA expression levels of a common anti-inflammatory cytokine, the *Il4,* to assess whether the liver microenvironment was more immunosuppressive in the absence of iNKT cells compared to *WT DEN* mice. Interestingly, we observed that the expression levels of *Il4* were significantly increased in *CD1d^−/−^ DEN* mice compared to *WT DEN* mice (Suppl. Fig. 4C). Furthermore, employing flow cytometry to analyze the immune response between *WT DEN* and *CD1d^−/−^ DEN* mice, we found that the ratio of T_regs_-to-T cells demonstrated a significant increase in the absence of iNKT cells compared to *WT DEN* mice in HCC (Suppl. Fig. 4D, E). Hence, the absence of iNKT cells appears to contribute to the generation of an immunosuppressive environment and enhanced cell proliferation following tumor establishment.

### iNKT cells promoted initiation of HCC

Because *CD1d^−/−^* mice developed fewer liver tumors, although with the same or even larger size, we hypothesized that iNKT cells may be required for HCC initiation. To test this hypothesis, we examined HCC development at various intervals (5, 7, and 9 months) after DEN treatment in *WT, Vα14^Tg/+^* and *CD1d^−/−^* mice (Figure 5A). At 5 months, *WT DEN* mice developed a few foci without any tumor development, whereas mice that lacked iNKT cells developed no foci or tumors (Figure 5B, upper panel, and C). Progressing to 7 months, *WT DEN* mice displayed both foci and small tumors, whereas *CD1d^−/−^ DEN* mice still showed no foci or tumors (Figure 5B middle panel, and C). In sharp contrast, *Vα14^Tg/+^* mice displayed significantly more foci than *WT* mice at 5 months after DEN treatment, accompanied by an equal number of small tumors at 7 months (figure 5B and C). By 9 months, both *WT DEN* and *CD1d^−/−^ DEN* mice developed HCC, as expected. However, mice lacking iNKT cells displayed significantly fewer foci and tumors compared to *WT* mice (Figure 5B lower panel, and C). Consistent with the previous time points, *Vα14^Tg/+^* mice showed a comparable phenotype to *WT* mice 9 months after DEN treatment. Taken together, our results show that the increased number of iNKT cells accelerated HCC development, while the absence of iNKT cells resulted in a delay in HCC development.

**Figure 5.**
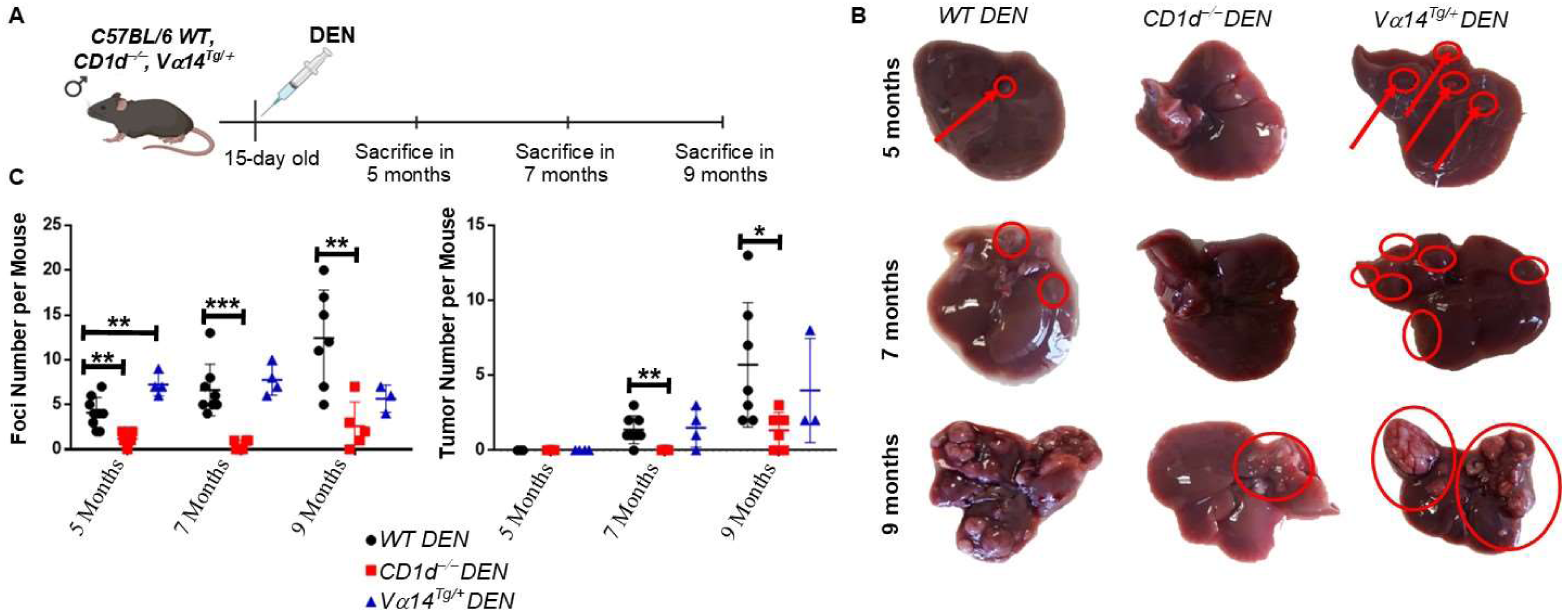
iNKT cells promoted initiation of HCC. **A** Graphical illustration of the experimental procedure. **B** Liver images from WT DEN, CD1d^−/−^ DEN and Vα14^Tg/+^ DEN mice in 5, 7 and 9 months after DEN treatment. **Red arrows:** foci. **C** Scatter plots show the foci and tumor numbers during HCC development in WT DEN, CD1d^−/−^ DEN and Vα14^Tg/+^ DEN mice. Bars represent mean ± SD from 3 independent experiments, with 1-3 mice in each time point. *, p<0.05, **, p<0.01 and ***, p<0.001 by t test.

### The lack of iNKT cells attenuated liver damage during initiation of HCC

Our results suggest that iNKT cells are required for the initiation of HCC after DEN treatment in mice. Therefore, we investigated the function of iNKT cells during acute inflammation 48 hours after DEN treatment in 15 days old mice. Histological analysis showed several damaged lesions in DEN-treated livers (Figure 7A), while blood alanine aminotransferase (ALT) and aspartate aminotransferase (AST) levels were elevated, indicating mild liver damage upon short-term DEN treatment (Figure 7B). The percentage and number of both iNKT and conventional T cells were reduced in DEN-treated *WT* mice, compared to non-treated mice (Figure 7C and D). Interestingly, iNKT cells showed an activated phenotype, assessed by the shift of CD69 expression (Figure 7E and F), while conventional T cells did not show signs of activation (Figure 7G).

**Figure 7.**
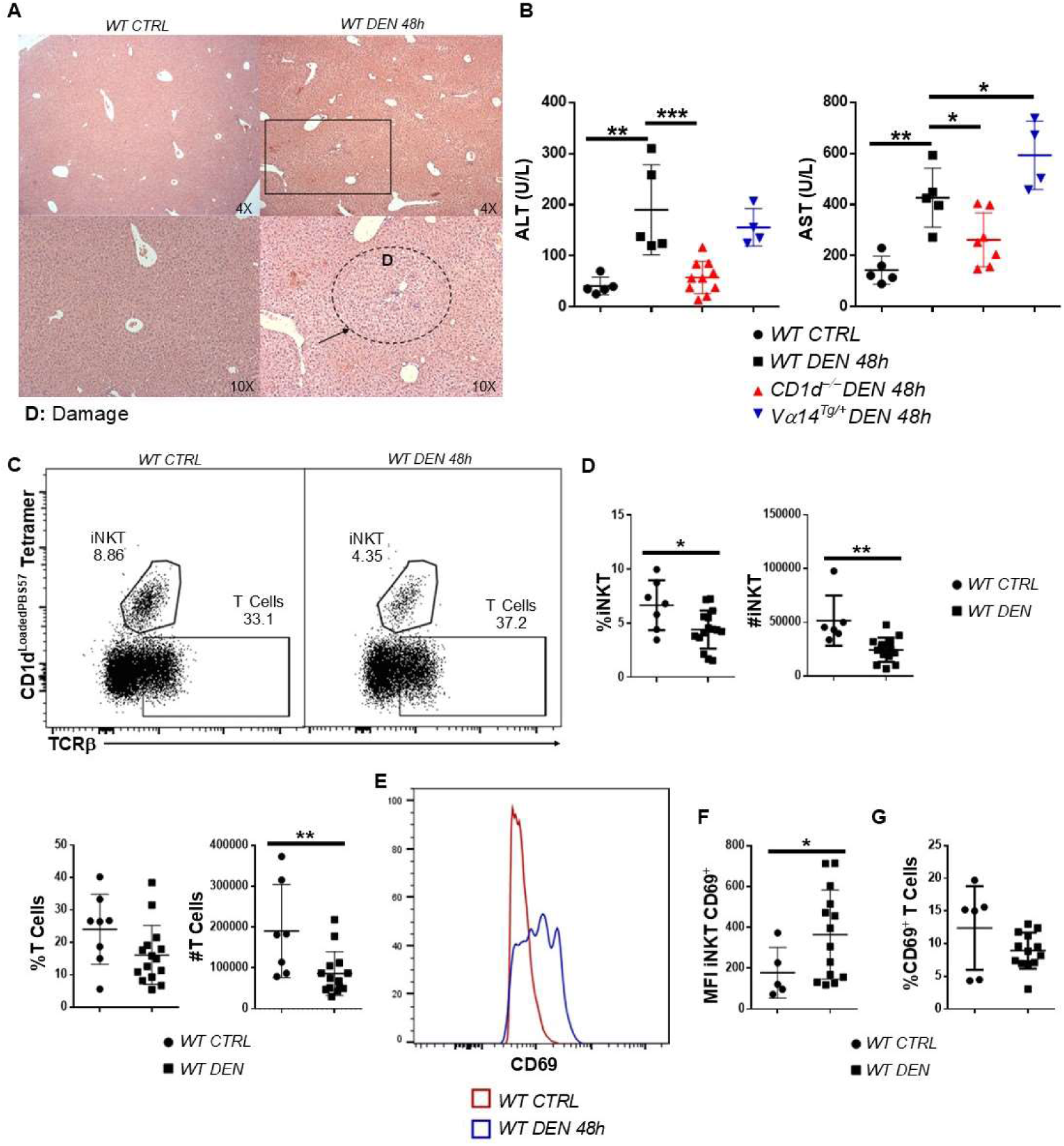
The lack of iNKT cells attenuated liver damage during initiation of HCC. **A** H&E histology from WT CTRL and WT DEN 48 hours after DEN injection in 4X and 10X magnification. **Black Arrow**, damaged lesions. **B** Scatter plots show the levels of liver transaminases ALT and AST from WT CTRL, WT DEN, CD1d^−/−^ DEN, and Vα14^Tg/+^ DEN mice 48h after DEN administration. **C** FACS plots show the frequency of iNKT and T cells in WT CTRL and WT DEN 48h mice. **D** Scatter plots show the percentage and number of iNKT and T cells. **E** Histogram shows the expression of CD69 in iNKT cells from WT CTRL and WT DEN 48h mice. **F** Scatter plot shows the MFI of CD69 in iNKT cells. **G** Scatter plot shows the expression of CD69 in T cells from WT CTRL and WT DEN 48h after DEN. Bars represent mean ± SD from 5 independent experiments. *, p<0.05, **, p<0.01, ***, p<0.001 by t test.

Remarkably, *CD1d^−/−^* DEN mice, which lack iNKT cells, displayed significantly lower ALT and AST levels compared to *WT DEN* mice, indicating reduced liver damage (Figure 7B). Conversely, *Vα14^Tg/+^* mice demonstrated significantly elevated AST levels compared to *WT DEN* mice, indicating the augmented degree of liver damage (Figure 7B).

In our effort to gauge inflammation levels, we quantified the expression of liver cytokines *Il1β*, *Il6*, and *TNFα,* which are indicators of inflammation, with real-time PCR. The mRNA levels of *Il1β*, *Il6*, and *TNFα* in the liver were markedly reduced in *CD1d^−/−^* DEN mice compared to *WT* DEN mice (Figure 8A), suggesting absence of inflammation in mice lacking iNKT cells. Consistent with this, flow cytometry analysis during acute inflammation revealed that *CD1d^−/−^ DEN* mice exhibited significantly lower percentage and number of neutrophils (CD11b^+^Ly6G^+^) compared to *WT DEN* mice (Figure 8B). In contrast, the frequency and number of neutrophils were unaffected in *Vα14^Tg/+^ DEN* compared to *WT DEN* mice (Suppl. Fig. 5A, B). Additionally, the percentage and activation^40,41^ of T_reg_ cells (TCRβ^+^CD4^+^Foxp3^+^) were notably increased in *CD1d^−/−^ DEN* mice compared to *WT DEN* mice (Figure 8C, D), whereas the frequency and activation of T_regs_ were significantly lower in *Vα14^Tg/+^ DEN* mice (Suppl. Fig. 5C, D, E). Collectively, these results indicate that iNKT cells promoted DEN-induced liver inflammation, which may in turn progress to HCC in mice.

**Figure 8.**
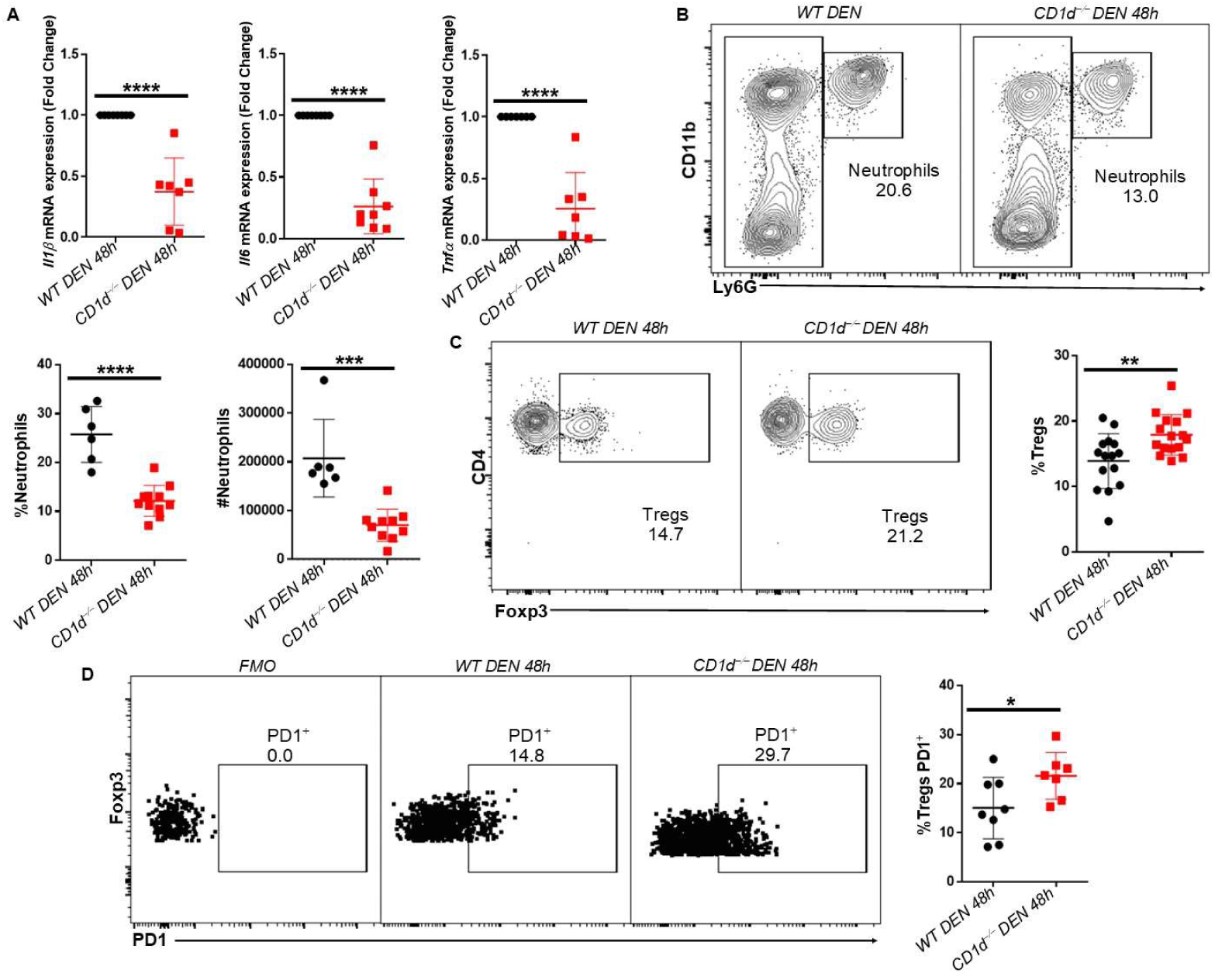
The lack of iNKT cells attenuated liver damage during initiation of HCC. **A** Scatter plots show the mRNA expression of the inflammatory markers Il1β, Il6, and TNFα in WT DEN and CD1d^−/−^ DEN 48h after DEN injection. **B** FACS plots of CD11b^+^Ly6G^+^ and scatter plots show the percentage and number of neutrophils. **C** FACS plots and scatter plot show TCRβ^+^CD4^+^Foxp3^+^ lymphocyte population **D** FACS plots and scatter plot show the percentage of activated (PD1^+^) Tregs. Bars represent mean ± SD from 5 independent experiments. *, p<0.05, **, p<0.01, ***, p<0.001, ****,

### Adoptive transfer of iNKT cells may suppress liver carcinogenesis

Because iNKT cells were reduced and dysfunctional in HCC livers, we hypothesized that provision of exogenous iNKT cells in established tumors may help tumor control. To examine this hypothesis, we adoptively transferred FACS-sorted iNKT cells from healthy mice into mice with HCC before and after tumor establishment. We injected iNKT cells intravenously at 1, 3, and 6 months after DEN treatment (*WT DEN* 1m, 3m, and 6m, respectively) and mice were euthanized 9 months after DEN treatment (Figure 9A and Suppl. Fig. 6).

**Figure 9.**
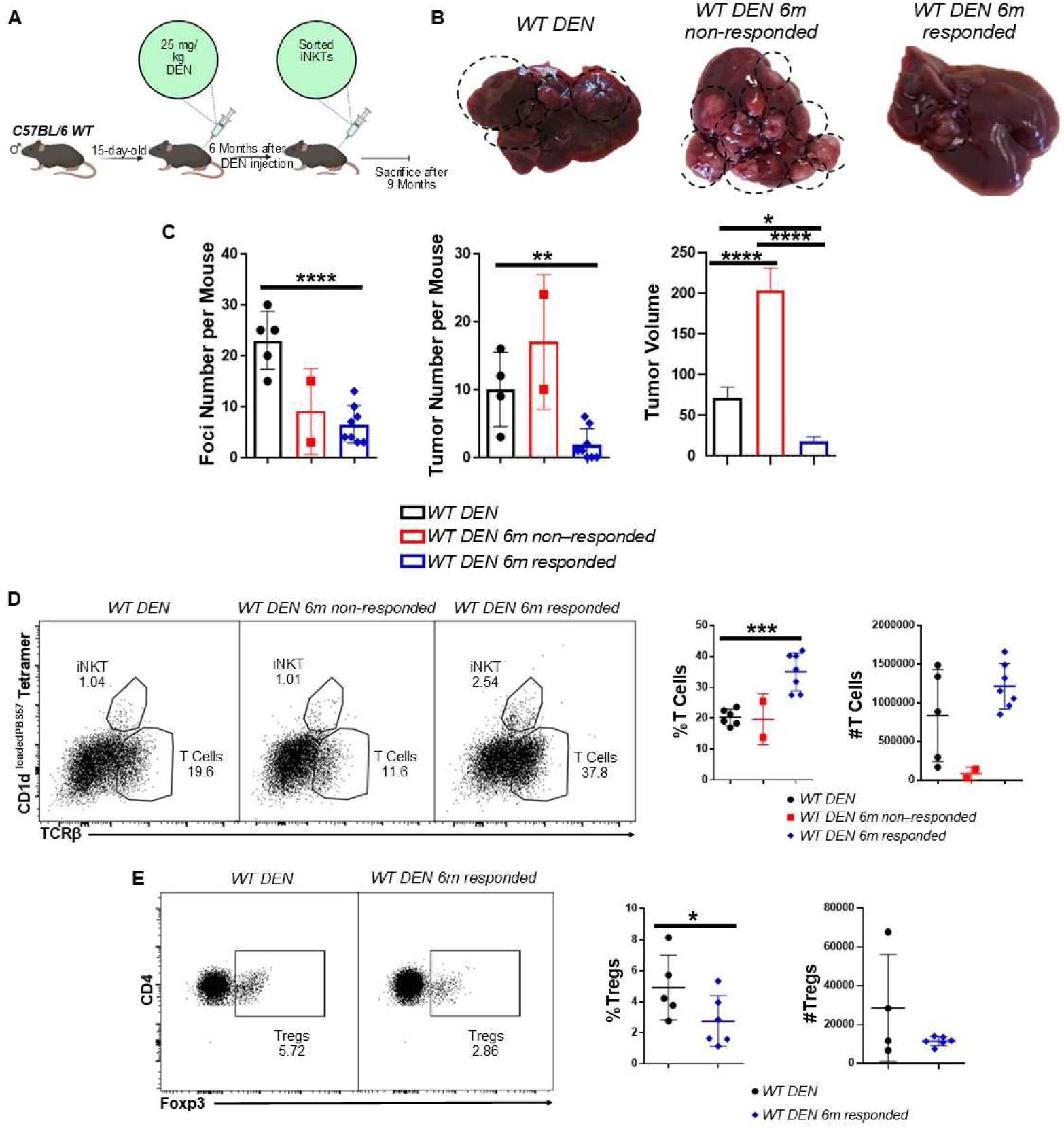
Adoptive transfer of iNKT cells may suppress liver carcinogenesis. **A** Graphical illustration of the experimental procedure. **B** Liver images from WT DEN, WT DEN that did not respond to iNKT cell adoptive transfer (WT DEN 6m non-responded), and WT DEN that respond to adoptive transfer of iNKT cells (WT DEN 6m responded) 9 months after DEN treatment. **C** Scatter plots show the foci and tumor numbers and bar plot shows the tumor volume. **D** FACS plots show the percentage of iNKT and T cells and scatter plots show the frequency and number of T cells in WT DEN, WT DEN 6m non-responded, and WT DEN 6m responded mice. **E** FACS plots show TCRβ^+^CD4^+^Foxp3^+^ lymphocyte population and scatter plots show their frequency and number in WT DEN and WT DEN 6m responded mice. Bars represent mean ± SD from 5 independent experiments, with one WT DEN and one or two WT DEN with iNKT cell adoptive transfer mice. *, p<0.05, **, p<0.01, ***, p<0.001, and ****, p<0.0001 by t test.

While *WT DEN* mice without iNKT adoptive transfer developed HCC as expected (Figure 9B), the majority (8/10 mice) of *WT DEN* mice with iNKT cell adoptive transfer 6 months post-DEN injection showed better tumor control, as shown by the significantly reduced number and volume of foci and tumors (Figure 9B and C). However, 2/10 *WT DEN 6m* mice did not respond effectively to iNKT cell treatment and developed HCC to a comparable level as *WT DEN* mice without adoptive transfer (Figure 9B, C). Flow cytometry analysis of these mice revealed that the frequency of iNKT cells was lower compared to *WT DEN 6m* mice that responded (Figure 9D), possibly indicating that iNKT cell transfer was not optimal. Nonetheless, these results suggest that iNKT cells may help to control tumor growth. Notably, *WT DEN 1m* and *3m* mice developed HCC to a similar extent as *WT DEN* mice that did not receive iNKT cells (Suppl. Fig. 6Α, Β), corroborating our previous findings showing that iNKT cells may promote early HCC development.

Furthermore, flow cytometry analysis of immune response in these mice revealed an increased frequency and number of T cells in *WT DEN* mice with iNKT adoptive transfer compared to *WT DEN* mice without adoptive transfer of iNKT cells (Figure 9D). *WT DEN* mice that did not respond to the adoptive transfer of iNKT cells have comparable frequency and number of T cells with *WT DEN* mice without adoptive transfer of iNKT cells (Figure 9D). Additionally, mice with iNKT cell adoptive transfer exhibited significantly decreased frequency of T_reg_ cells compared to *WT DEN* mice without iNKT cell adoptive transfer (Figure 9E). Taken together, our findings suggest that iNKT cells may act therapeutically to inhibit progression of established HCC.

## Discussion

In this study, we aimed to elucidate the role of iNKT cells in the development and progression of HCC. Our research revealed a significant reduction in the iNKT cell population and a loss of their functionality within HCC, resulting in decreased activation, increased exhaustion, and impaired cytokine production. Additionally, iNKT cells underwent apoptosis within the HCC microenvironment, leading to their reduced numbers. *CD1d*-deficient mice, which lack iNKT cells, developed significantly fewer number of foci and tumors, albeit with larger volume. Lack of iNKT cells contributed also to enhanced immunosuppression, marked by elevated frequency of T_reg_ cells and increased cell proliferation after tumor establishment. Furthermore, iNKT cells mitigated liver damage during HCC initiation, as young mice lacking iNKT cells showed reduced DEN-mediated inflammation and liver damage. Notably, adoptive transfer of healthy iNKT cells suppressed liver carcinogenesis after tumors have been established. These findings indicate that iNKT cells have a dual role in HCC, both promoting the initiation and development of HCC while also combating its growth and progression.

Our research showed a significant reduction in both the frequency and number of iNKT cells in HCC, in contrast to conventional T cells, which were increased in numbers. Importantly, the remaining tumoral iNKT cells showed signs of dysfunction, which is in accordance with the phenotype of human iNKT cells reported in HCC patients^42,43^. Similar to iNKT cells, the number of hepatic MAIT cells was also reduced in HCC patients^44,45^, indicating a global loss of T cells with innate characteristics from the HCC tumor microenvironment. The tumor itself may directly promote lymphocyte apoptosis through Fas/FasL interactions with T cells^46^. Notably, through RNA sequencing and flow cytometry for active caspase 3 detection, we found that iNKT cells in HCC were apoptotic, which may account for their reduction – an observation not previously reported. The dysfunctional state of iNKT cells may be induced by continuous TCR stimulation by tumor antigens and/or chronic inflammation. Indeed, iNKT cells become anergic or immunosuppressive after repeated α-GalCer stimulation^47,48^, while a recent study indicated that long-chain acylcarnitines derived from HCC tumor cells led to premature iNKT cell senescence and impaired α-GalCer responsiveness^43^. Although the potential antigens that may chronically and directly stimulate iNKT cells to exhaustion during HCC are yet to be defined, our data suggest that iNKT cell loss and/or dysfunction is an important characteristic of the HCC immune microenvironment.

Our study demonstrated that the lack of iNKT cells in *CD1d^−/−^ DEN* mice ameliorated HCC development, as shown by the reduced tumor and foci number observed in these mice. In addition, our kinetic analysis determined that *CD1d^−/−^* developed precancerous foci at slower pace than their *WT* counterparts, possibly indicating that the lack of iNKT cells delayed the initiation of tumorigenesis. Two additional lines of evidence support this hypothesis. First, adoptive transfer of healthy iNKT cells into *WT DEN* mice, conducted 1- and 3-months post-DEN injection, revealed that these mice developed HCC to a similar extent to, if not more severe than, *WT DEN* mice that did not receive iNKT cells. Consistent with that, *Vα14^Tg/+^* mice, which have enhanced development of iNKT cells and thus excess iNKT numbers early in life, showed accelerated HCC tumorigenesis, evidenced by increased numbers of foci at the 5-month time point compared to *WT* mice. Therefore, iNKT cell activity was detrimental during the early stages of HCC tumorigenesis.

In support of this hypothesis, 15-day old *CD1d-*deficient mice treated short-term with DEN for 48 hours exhibited lower hepatic damage and acute inflammation, whereas *Vα14^Tg/+^ DEN* mice showed increased liver damage compared to *WT DEN* mice. Importantly, iNKT cells upregulated their surface CD69 expression upon short-term DEN treatment, indicating that they mediate, and may potentially perpetuate, DEN-induced acute liver inflammation. These results are in accordance with the deleterious role of activated iNKT cells in promoting hepatitis upon α-GalCer or LPS treatment^49,50^. Moreover, in the absence of iNKT cells, the T_reg_ population and their activation significantly increased, potentially facilitating a faster resolution of acute inflammation, and protecting *CD1d^−/−^ DEN* mice from chronic liver damage. In contrast, mice with overproduction of iNKT cells showed significantly lower frequency, number, and activation of T_reg_ cells during short-term DEN treatment, suggesting an inability of these mice to resolve acute inflammation, thus resulting in enhanced liver damage. Our results are consistent with previous results showing an inverse correlation between the number of T_reg_ cells and the development of liver inflammation and damage in various mouse models of liver disease^51,52,53^.

Additionally, activated iNKT cells have been correlated with decreased T_reg_ cell percentage and production of the immunosuppressant cytokines IL10 and TGFβ^54^. Although the underlying mechanisms remain to be elucidated, the interaction between iNKT and T_reg_ cells may regulate chronic liver inflammation during HCC development. Additionally, independent studies have shown that iNKT cells enhanced lipid uptake from hepatocytes, contributing to NASH and HCC development in mice fed with Choline Deficient-High Fat Diet (CD-HFD)^29^. Regardless of the mechanisms involved, our results suggest that iNKT cells promoted the initiation of HCC, through perpetuation of acute inflammation and enhanced liver damage.

Despite the decreased number of foci and tumors in DEN-treated *CD1d^−/−^* mice, we noted that the size of the few developed tumors significantly increased compared to *WT* DEN mice, whereas it was significantly reduced in *Vα14^Tg/+^ DEN* mice. Tumors that grew in the absence of iNKT cells showed increased proliferation, as evidenced by higher expression of the cell cycle marker Ki67 and the proliferation marker *c-Jun.* Furthermore, we observed an expansion of T_reg_ cells in the absence of iNKT cells during the late stages of HCC, which contributed to an enhanced immunosuppressive microenvironment, exemplified by the augmented expression of *Il4*. The reduction in the number of T_reg_ cells in TGF-β1 knockout mice led to decreased metastasis in mice with orthotopic HCC^55^, while secretion of immunosuppressive cytokines by T_regs_ contributed to impaired cancer immunosurveillance^52,56^.These results collectively suggest that iNKT cells may effectively control HCC tumor growth, possibly through interactions with T_reg_ cells. In accordance with this hypothesis, adoptive transfer of iNKT cells, isolated *ex vivo* from healthy mice, into *WT* mice with established HCC led to reduced numbers and shrinkage of HCC tumors, accompanied by reduced T_reg_ cell frequency in eight out of ten treated mice, while two mice showed no improvement. We postulate that this heterogeneous response may be due to the variable stage of HCC development at the time of iNKT cell transfer. Nonetheless, our results showcase the potential of iNKT cells to be used therapeutically for the treatment against HCC. Indeed, adoptive transfer of *in vitro* expanded autologous iNKT cells in HCC patients reduced α-fetoprotein (AFP) levels and improved overall and progression-free survival in four out of ten patients with HCC, while one patient showed no tumor recurrence^57^. Taken together, our data support that iNKT cells play a dual role in HCC development and progression: first, they promote acute inflammation and subsequent chronic liver damage leading to HCC. Second, once HCC tumors are established, iNKT cells help to control tumor growth and HCC progression.

## Material and Methods

### Mice

Mice of the *C57BL/6* background were used in this study, including *WT*, *CD1d^−/−^*, and *Vα14^Tg/+^* mice. *CD1d^−/−^* and *Vα14^Tg/+^* mice have been previously described. 15-day-old male mice were injected i.p. with 25mg/kg diethyl- nitrosamine (DEN) (Sigma, Cat. #: N0756-10ML) to induce HCC and mice were sacrificed 38-42 weeks later, or at the indicated time points. For DEN-induced acute inflammation, 100 mg/kg DEN were injected to 15-day-old male mice and mice were euthanized 48 hours later.

### Quantifications

To determine the %Ratio Liver/Body weight, we used the formula: %Ratio = (Liver Weight (g) / Body Weight (g)) x 100 for each mouse. Additionally, tumors were defined as cancer lesions larger than 3 mm, while foci were defined as lesions smaller than 3 mm. To calculate liver size, we applied the following formula, for each liver tumor in every mouse: Tumor Volume = 0.5 x L x W^2, where L represents the tumor length and W represents the tumor width. The lymphocyte number was calculated as #Lymphocytes = %Lymphocytes x %Live Lymphocytes x #Total cells considering the total cell count measured using the Neubauer chamber. The count of CD45^+^ cells was computed as follows: #CD45^+^ = #lymphocytes x %CD45 ^+^.

### Histology (H&E staining)

Formalin (3 ml) was added into liver samples from the indicated mouse strains. Subsequently, paraffin was incorporated into livers using histokinette, and paraffin blocks were created. The tissue was then sectioned into 4 μm segments, and slides were stained with hematoxylin and eosin.

### RNA extraction and Real-Time PCR

Total liver RNA was isolated from homogenized liver tissue with TRIzol (PanReac AppliChem, Cat. #: A4051,0100) according to the manufacturer’s instructions. For sorted cells, RNA was extracted using RNAeasy micro kit (Qiagen, Cat. #: 74034). Reverse transcription was performed using Superscript III (Invitrogen, Cat. #: 18080-044). Real-Time PCR was conducted with gene-specific primers in STEPOne and LightCycle96 Roche, using the FAST START Universal SYBR Green Master (Roche, Cat. #: 4913914001). *Gapdh* served as reference gene. The relative expression was calculated using the Δ^CT^ method. The following sets of primers were used: ***Gapdh***, Forward (Fw): 5’GTTGTCTCCTGCGACTTCA3’ and Reverse (Rv): 5’GGTGGTCCAGGGTTTCTTA3’, ***V****α****14J****α****18***, *Vα14*: 5’TCCTCAGTCCCTGGTTGTC3’ And *Jα18*: 5’CAAAATGCAGCCTCCCTAAG3’, ***Afp***, Forward (Fw): 5’CTCAGCGAGGAGAAATGGTC3’ and Reverse (Rv): 5’GAGTTCACAGGGCTTGCTTC3’, ***Gpc3***, 5’TGAGCCGGTGGTTAGCCAG3’ and Reverse (Rv): 5’CTTCACTTTCACCATCCCGTC3’, ***c-Jun***, Forward (Fw): 5’CCTTCTACGACGATGCCTTC3’ and Reverse (Rv): 5’GGCCAGGTTCAA GGTCATGC3’, ***Il4***, Forward (Fw): 5’ACTCTAGTGTTCTCATGGAGC3’ and Reverse (Rv): 5’TTCAGTGATGTGGACTTGGAC3’, ***Il1****β*, Forward (Fw): 5’GCAACTGTTCCTGAACTCAACT3’ and Reverse (Rv): 5’GGTCCGTCAACTTCAAAGAAC3’, ***Il6***, Forward (Fw): 5’CTGCAAGAGACTTCCATCCAG3’ and Reverse (Rv): 5’GTGGTATAGACAGGTCTGTTGG3’, ***Tnf****α*, Forward (Fw): 5’GGCCTCCCTCTCATCAGTTC3’ and Reverse (Rv): 5’GGTGGTTTGCTACGACGTGG3’. Lymphocyte Isolation To isolate hepatic lymphocytes, livers were placed in a 100 μm filter and homogenized using FACS buffer (HBSS, 0.25% BSA, 25 mg/ml DNase I, 0.1% NaN_3_). Subsequently, samples were centrifuged at 1500 rpm for 5 min at 4°C, and pellets were resuspended in 8 ml Percoll mix (for 10 Percoll mix: 0.4 ml 10X PBS, 4 ml Percoll, 5.6 ml 1X PBS). Samples were centrifuged at 2000 rpm for 20 min at 20°C without braking. Red blood cells were lysed with H_2_O and lymphocytes were resuspended in 1 ml FACS Buffer after centrifugation at 1500 rpm, 5 min, 4°C.

### Flow cytometry, Cell Sorting and Antibodies

Samples were acquired using FACSCanto III or FACSCelesta, or were sorted with FACSAria III, and then analyzed using FlowJo v10. Lymphocytes were incubated with antibodies for 30 min in ice and in dark, in the presence of anti-mouse CD16/32 Fc block (eBioscience, Cat #: 14-0161-85) and Propidium Iodide (eBioscience, Cat #: 00-6990-42). The following antibodies were purchased from eBioscience and Biolegend: TCRβ (H57-597), CD4 (GK1.5), CD8a (53-6.7), CD69 (H1.2F3), PD1 (RMPI-30), CD11b (M1/70), F4/80 (BM8), Ly6G (1A8), CD206 (C068C2), CD80 (16-10A1), IFNγ (MG1.2), TNFα (MP6- XT22), CD45 (30-F11), Foxp3 (150D), NK1.1 (PK136). Detection of iNKT cells was performed with CD1d^LoadedPBS^^57^ Tetramers provided by the NIH tetramer facility at Emory University. iNKT cells were sorted as CD1d^Loaded PBS^^57^ Tetramer^+^ TCRβ^+^ cells.

To conduct intracellular staining for the transcription factor Foxp3, we used the Foxp3/Transcription Factor Staining Buffer Set (eBioscience, Cat #: 00-5523-00). The Pyroptosis/Caspase 1 assay kit, green (ImmunoChemistry Technologies, Cat #: SKU: 9145) and the Fam-Flica Caspase 3&7 assay kit (ImmunoChemistry Technologies, Cat #: SKU: 93) were used for staining for active caspase-1 and active caspase-3, respectively. These assays were conducted following the manufacturer’s instructions.

### Cell culture, Cytokine Production and Staining

Lymphocytes were exposed to 500 ng/ul phorbol 12-myristate 13- acetate (PMA), 1 μg/ul ionomycin (ION), and Brefeldin A for 4 hours, before surface staining with CD1d^Loaded PBS^^57^ Tetramer and TCRβ, followed by intracellular staining for IFNγ and TNFα using Intracellular Fixation & Permeabilization Buffer Set (eBioscience, Cat. # 88-8824-00).

### Immunofluorescence

Liver tissue sections were deparaffinized and hydrated using the following sequence of reagents, each for 5 min: Xylene (3 times), 100% Ethanol (2 times), 90% Ethanol, 70% Ethanol, 50% Ethanol, 30% Ethanol, and ddH2O (2 times). Slides were then inserted into pre-heated Citrate Buffer (90°C) for 30 minutes and washed with ddH_2_O for 2 min, twice. Subsequently, liver sections were fixed using 4% paraformaldehyde (PFA) for 15 min and washed with 1X PBS for 5 min. A hydrophobic circle around the liver tissue was created using PAPpen (Kisker), and blocking was performed using 10% BSA in 0.1% TritonX/PBS. Liver sections were then stained with Ki67 monoclonal antibody (eBioscience, SP6, Cat. #: MA5-14520) (diluted 1/100) overnight at 4°C. After washing the sections with 1X PBS for 5 min (3 times), liver sections were stained with goat anti-Rabbit IgG (H+L) Cross-Adsorbed Secondary Antibody, Alexa Fluor 647 (Thermo Fisher Scientific, Cat. #: A21244) (diluted 1/500) for 3 hours at room temperature. Finally, coverslips were mounted onto slides using DAPI Mounting Medium (Ibidi, Cat. #: 50011). Images were acquired using a TCS SP8X confocal system (Leica), and image analysis was performed using Fiji-ImageJ.

### RNA sequencing and Data Analysis

Libraries were prepared using the 3′ mRNA-Seq Library Prep Kit Protocol for Ion Torrent (QuantSeq-LEXOGEN™ Vienna, Austria), according to manufacturer’s instructions. Briefly, up to 500 ng of sample RNA were used for first strand synthesis, followed by removal of RNA and initiation of second strand synthesis by a random primer containing Ion Torrent compatible linker sequences. Barcodes were introduced and second-strand synthesis was performed, followed by magnetic bead-based purification, amplification of the resulting libraries and re-purification. The libraries were assessed for quality and quantity on a bioanalyzer using the DNA High Sensitivity Kit reagents and protocol (Agilent Technologies). The quantified libraries were pooled and templated and enriched on the Ion Proton One Touch system using the Ion PI™ Hi-Q™ OT2 200 Kit (ThermoFisher Scientific); they were sequenced on Ion Proton PI™ V2 chips using the Ion PI™ Hi-Q™ Sequencing 200 Kit (ThermoFisher Scientific) on an Ion Proton™ System, according to manufacturer’s instructions.

Fastq files were mapped to mouse genomic build mm10 using a two-step alignment process. In the first round, reads were mapped with hisat2^58^ using the default parameters and then, the unmapped reads were mapped with bowtie2^59^ using the --local and --very-sensitive parameters. RNA-seq analysis was conducted with the Bioconductor package metaseqR2^60^, as previously described^61^. Raw reds were estimated through metaseqR2 and were normalized with the DESEQ algorithm. Differential gene expression was conducted with the PANDORA algorithm^62^ by combining the algorithms DESeq2^63^, edgeR^64^, limma^65^, NBPSeq^66^, and NOISeq^67^. Genes exhibiting a meta-p-value < 0.05 and a fold change greater than 1.0 or less than −1.0 on the log2 scale (corresponding to a 2-fold change in the natural scale) were considered differentially expressed. The volcano plot was generated using ggplot2. The code for the analysis is available in https://github.com/alex-galaras/verykokakis_FFDEN_analysis.git. ShinyGo analysis (ShinyGo v0.741) was used to uncover Gene Ontology (GO) classifications related to biological processes.

### Biochemical Analysis

Blood serum was obtained from mice through two consecutive centrifugation steps at 3000 rpm and 6000 rpm for 10 min at 20°C. Following centrifugation, the serum was divided into aliquots of 45 μl for each sample. Each sample was then diluted 1:1 with ddH_2_O. The levels of liver transaminases ALT and AST, were subsequently measured using a biochemical analyzer.

### Adoptive Transfer of iNKT cells

Livers were isolated and pooled from 3-5 *WT* mice and passed through a 100 μm cell strainer. Following centrifugation, samples were resuspended in a Percoll mix and centrifuged without braking. Subsequently, red blood cell lysis was performed, and lymphocytes were finally resuspended in MACS Buffer (HBSS, 0.25% BSA, and 25mg/ml DNase I). Lymphocytes were then stained with CD1d^LoadedPBS^^57^ Tetramer and anti-TCRβ for iNKT cell detection. Propidium Iodide (PI) was utilized to exclude dead cells. iNKT cells were sorted as CD1d^Loaded PBS^^57^ Tetramer^+^TCRβ^+^, using FACSAria III. For adoptive transfer, iNKT cells were resuspended in 2.5% FBS/PBS, and 1x10^5^ iNKT cells were intravenously injected into each recipient.

### Statistical Analysis

Statistical analysis was conducted using t-test analysis in GraphPad Prism 6 and 8. *, p<0.05, **, p<0.01, ***, p<0.001, ****, p<0.0001.

## Acknowledgements

The authors would like to thank members of the Verykokakis, Fousteri, and Hatzis labs for helpful discussions and for providing reagents, Manos Aerakis and Mariela Alvanou for their insightful discussions and contributions, and the NIH Tetramer Facility for providing CD1d^LoadedPBS57^ tetramers for detection of iNKT cells. The authors are grateful to the Flow Cytometry Facility, the Histology Facility, and the Genomics Facility at BSRC Alexander Fleming for contributions to this work. This work was benefited by HFRI research grants No 486 and 14672 to MV. MP was supported by the 4^th^ Call for HFRI PhD fellowships No 11115. TR-T was supported by “Nuclear Receptor-Network” Consortium Grant PITN-GA-2013-606806 from the European Union 7^th^ Framework Programme Marie Curie Initial Training Network and as part of the PEOPLE-2013 program. We acknowledge support of this work by Research Infrastructure InfrafrontierGR (MIS 5002802) funded by the Operational Programme “Competitiveness, Entrepreneurship and Innovation” (NSRF 2014- 2020) and by project MIS 6004752 funded by the Regional Operational Programme “ATTICA” (NSRF 2021-2027), both of which were co-financed by Greece and the European Union (European Regional Development Fund).

## References

1. Trefts E, Gannon M, Wasserman DH. The liver. Curr Biol. Nov 06 2017;27(21):R1147- R1151. doi:10.1016/j.cub.2017.09.019

2. Chen D, Le TH, Shahidipour H, Read SA, Ahlenstiel G. The Role of Gut-Derived Microbial Antigens on Liver Fibrosis Initiation and Progression. Cells. 10 27 2019;8(11)doi:10.3390/cells8111324

3. Ficht X, Iannacone M. Immune surveillance of the liver by T cells. Sci Immunol. 09 04 2020;5(51)doi:10.1126/sciimmunol.aba2351

4. Siegel RL, Miller KD, Fuchs HE, Jemal A. Cancer statistics, 2022. CA Cancer J Clin. 01 2022;72(1):7-33. doi:10.3322/caac.21708

5. Fu J, Xu D, Liu Z, et al. Increased regulatory T cells correlate with CD8 T-cell impairment and poor survival in hepatocellular carcinoma patients. Gastroenterology. Jun 2007;132(7):2328–39. doi:10.1053/j.gastro.2007.03.102

6. Ninomiya T, Akbar SM, Masumoto T, Horiike N, Onji M. Dendritic cells with immature phenotype and defective function in the peripheral blood from patients with hepatocellular carcinoma. J Hepatol. Aug 1999;31(2):323–31. doi:10.1016/s0168-8278(99)80231-1

7. Wu K, Kryczek I, Chen L, Zou W, Welling TH. Kupffer cell suppression of CD8+ T cells in human hepatocellular carcinoma is mediated by B7-H1/programmed death-1 interactions. Cancer Res. Oct 15 2009;69(20):8067–75. doi:10.1158/0008-5472.CAN-09-0901

8. Lu LC, Chang CJ, Hsu CH. Targeting myeloid-derived suppressor cells in the treatment of hepatocellular carcinoma: current state and future perspectives. J Hepatocell Carcinoma. 2019;6:71–84. doi:10.2147/JHC.S159693

9. Ghiringhelli F, Ménard C, Martin F, Zitvogel L. The role of regulatory T cells in the control of natural killer cells: relevance during tumor progression. Immunol Rev. Dec 2006;214:229–38. doi:10.1111/j.1600-065X.2006.00445.x

10. Zahran AM, Abdel-Meguid MM, Ashmawy AM, et al. Frequency and Implications of Natural Killer and Natural Killer T Cells in Hepatocellular Carcinoma. Egypt J Immunol. Jun 2018;25(2):45–52.

11. Egawa T, Eberl G, Taniuchi I, et al. Genetic evidence supporting selection of the Valpha14i NKT cell lineage from double-positive thymocyte precursors. Immunity. Jun 2005;22(6):705–16. doi:10.1016/j.immuni.2005.03.011

12. Gapin L, Matsuda JL, Surh CD, Kronenberg M. NKT cells derive from double-positive thymocytes that are positively selected by CD1d. Nat Immunol. Oct 2001;2(10):971–8. doi:10.1038/ni710

13. Lee YJ, Holzapfel KL, Zhu J, Jameson SC, Hogquist KA. Steady-state production of IL-4 modulates immunity in mouse strains and is determined by lineage diversity of iNKT cells. Nat Immunol. Nov 2013;14(11):1146–54. doi:10.1038/ni.2731

14. Wu L, Van Kaer L. Natural killer T cells in health and disease. Front Biosci (Schol Ed). 01 01 2011;3(1):236-51. doi:10.2741/s148

15. Cullen R, Germanov E, Shimaoka T, Johnston B. Enhanced tumor metastasis in response to blockade of the chemokine receptor CXCR6 is overcome by NKT cell activation. J Immunol. Nov 01 2009;183(9):5807–15. doi:10.4049/jimmunol.0803520

16. Coquet JM, Chakravarti S, Kyparissoudis K, et al. Diverse cytokine production by NKT cell subsets and identification of an IL-17-producing CD4-NK1.1- NKT cell population. Proc Natl Acad Sci U S A. Aug 12 2008;105(32):11287–92. doi:10.1073/pnas.0801631105

17. Karadimitris A, Ripoll-Fiol C, Guerra JC. Invariant NKT cells as a platform for CAR immunotherapy and prevention of acute Graft-versus-Host Disease. Hemasphere. Jun 2019;3(Suppl)doi:10.1097/HS9.0000000000000220

18. Haraguchi K, Takahashi T, Nakahara F, et al. CD1d expression level in tumor cells is an important determinant for anti-tumor immunity by natural killer T cells. Leuk Lymphoma. Oct 2006;47(10):2218–23. doi:10.1080/10428190600682688

19. Bassiri H, Das R, Guan P, et al. iNKT cell cytotoxic responses control T-lymphoma growth in vitro and in vivo. Cancer Immunol Res. Jan 2014;2(1):59–69. doi:10.1158/2326-6066.CIR-13-0104

20. Parekh VV, Lalani S, Van Kaer L. The in vivo response of invariant natural killer T cells to glycolipid antigens. Int Rev Immunol. 2007 Jan-Apr 2007;26(1-2):31-48. doi:10.1080/08830180601070179

21. Metelitsa LS. Anti-tumor potential of type-I NKT cells against CD1d-positive and CD1d- negative tumors in humans. Clin Immunol. Aug 2011;140(2):119–29. doi:10.1016/j.clim.2010.10.005

22. De Santo C, Salio M, Masri SH, et al. Invariant NKT cells reduce the immunosuppressive activity of influenza A virus-induced myeloid-derived suppressor cells in mice and humans. J Clin Invest. Dec 2008;118(12):4036–48. doi:10.1172/JCI36264

23. Shin T, Nakayama T, Akutsu Y, et al. Inhibition of tumor metastasis by adoptive transfer of IL-12-activated Valpha14 NKT cells. Int J Cancer. Feb 15 2001;91(4):523–8. doi:10.1002/1097-0215(20010215)91:4<523::aid-ijc1087>3.0.co;2-l

24. Cui J, Shin T, Kawano T, et al. Requirement for Valpha14 NKT cells in IL-12-mediated rejection of tumors. Science. Nov 28 1997;278(5343):1623-6. doi:10.1126/science.278.5343.1623

25. Nakagawa R, Motoki K, Ueno H, et al. Treatment of hepatic metastasis of the colon26 adenocarcinoma with an alpha-galactosylceramide, KRN7000. Cancer Res. Mar 15 1998;58(6):1202-7.

26. Heczey A, Liu D, Tian G, et al. Invariant NKT cells with chimeric antigen receptor provide a novel platform for safe and effective cancer immunotherapy. Blood. Oct 30 2014;124(18):2824–33. doi:10.1182/blood-2013-11-541235

27. Tian G, Courtney AN, Jena B, et al. CD62L+ NKT cells have prolonged persistence and antitumor activity in vivo. J Clin Invest. 06 01 2016;126(6):2341-55. doi:10.1172/JCI83476

28. Wehr A, Baeck C, Heymann F, et al. Chemokine receptor CXCR6-dependent hepatic NK T Cell accumulation promotes inflammation and liver fibrosis. J Immunol. May 15 2013;190(10):5226–36. doi:10.4049/jimmunol.1202909

29. Wolf MJ, Adili A, Piotrowitz K, et al. Metabolic activation of intrahepatic CD8+ T cells and NKT cells causes nonalcoholic steatohepatitis and liver cancer via cross-talk with hepatocytes. Cancer Cell. Oct 13 2014;26(4):549–64. doi:10.1016/j.ccell.2014.09.003

30. Mossanen JC, Tacke F. Role of lymphocytes in liver cancer. Oncoimmunology. Nov 01 2013;2(11):e26468. doi:10.4161/onci.26468

31. Papanastasatou M, Verykokakis M. Innate-like T lymphocytes in chronic liver disease. Front Immunol. 2023;14:1114605. doi:10.3389/fimmu.2023.1114605

32. Miyagi T, Takehara T, Tatsumi T, et al. CD1d-mediated stimulation of natural killer T cells selectively activates hepatic natural killer cells to eliminate experimentally disseminated hepatoma cells in murine liver. Int J Cancer. Aug 10 2003;106(1):81–9. doi:10.1002/ijc.11163

33. Margalit M, Shibolet O, Klein A, et al. Suppression of hepatocellular carcinoma by transplantation of ex-vivo immune-modulated NKT lymphocytes. Int J Cancer. Jun 20 2005;115(3):443–9. doi:10.1002/ijc.20889

34. Mossanen JC, Kohlhepp M, Wehr A, et al. CXCR6 Inhibits Hepatocarcinogenesis by Promoting Natural Killer T- and CD4. Gastroenterology. 05 2019;156(6):1877–1889.e4. doi:10.1053/j.gastro.2019.01.247

35. Anson M, Crain-Denoyelle AM, Baud V, et al. Oncogenic β-catenin triggers an inflammatory response that determines the aggressiveness of hepatocellular carcinoma in mice. J Clin Invest. Feb 2012;122(2):586–99. doi:10.1172/JCI43937

36. Vesselinovitch SD, Mihailovich N. Kinetics of diethylnitrosamine hepatocarcinogenesis in the infant mouse. Cancer Res. Sep 1983;43(9):4253–9.

37. Lee JS, Chu IS, Mikaelyan A, et al. Application of comparative functional genomics to identify best-fit mouse models to study human cancer. Nat Genet. Dec 2004;36(12):1306–11. doi:10.1038/ng1481

38. Exley MA, Bigley NJ, Cheng O, et al. Innate immune response to encephalomyocarditis virus infection mediated by CD1d. Immunology. Dec 2003;110(4):519–26. doi:10.1111/j.1365-2567.2003.01779.x

39. Griewank K, Borowski C, Rietdijk S, et al. Homotypic interactions mediated by Slamf1 and Slamf6 receptors control NKT cell lineage development. Immunity. Nov 2007;27(5):751–62. doi:10.1016/j.immuni.2007.08.020

40. McGrath MM, Najafian N. The role of coinhibitory signaling pathways in transplantation and tolerance. Front Immunol. 2012;3:47. doi:10.3389/fimmu.2012.00047

41. Jubel JM, Barbati ZR, Burger C, Wirtz DC, Schildberg FA. The Role of PD-1 in Acute and Chronic Infection. Front Immunol. 2020;11:487. doi:10.3389/fimmu.2020.00487

42. Tao L, Wang S, Kang G, et al. PD-1 blockade improves the anti-tumor potency of exhausted CD3. Oncoimmunology. 2021;10(1):2002068. doi:10.1080/2162402X.2021.2002068

43. Cheng X, Tan X, Wang W, et al. Long-Chain Acylcarnitines Induce Senescence of Invariant Natural Killer T Cells in Hepatocellular Carcinoma. Cancer Res. Feb 15 2023;83(4):582–594. doi:10.1158/0008-5472.CAN-22-2273

44. Huang W, Ye D, He W, He X, Shi X, Gao Y. Activated but impaired IFN-γ production of mucosal-associated invariant T cells in patients with hepatocellular carcinoma. J Immunother Cancer. 11 2021;9(11)doi:10.1136/jitc-2021-003685

45. Zheng C, Zheng L, Yoo JK, et al. Landscape of Infiltrating T Cells in Liver Cancer Revealed by Single-Cell Sequencing. Cell. Jun 15 2017;169(7):1342–1356.e16. doi:10.1016/j.cell.2017.05.035

46. Whiteside TL. Apoptosis of immune cells in the tumor microenvironment and peripheral circulation of patients with cancer: implications for immunotherapy. Vaccine. Dec 19 2002;20 Suppl 4:A46–51. doi:10.1016/s0264-410x(02)00387-0

47. Parekh VV, Wilson MT, Olivares-Villagómez D, et al. Glycolipid antigen induces long-term natural killer T cell anergy in mice. J Clin Invest. Sep 2005;115(9):2572–83. doi:10.1172/JCI24762

48. Sag D, Krause P, Hedrick CC, Kronenberg M, Wingender G. IL-10-producing NKT10 cells are a distinct regulatory invariant NKT cell subset. J Clin Invest. Sep 2014;124(9):3725–40. doi:10.1172/JCI72308

49. Biburger M, Tiegs G. Alpha-galactosylceramide-induced liver injury in mice is mediated by TNF-alpha but independent of Kupffer cells. J Immunol. Aug 01 2005;175(3):1540–50. doi:10.4049/jimmunol.175.3.1540

50. Nagarajan NA, Kronenberg M. Invariant NKT cells amplify the innate immune response to lipopolysaccharide. J Immunol. Mar 01 2007;178(5):2706–13. doi:10.4049/jimmunol.178.5.2706

51. Ikeno Y, Ohara D, Takeuchi Y, et al. Foxp3+ Regulatory T Cells Inhibit CCl4-Induced Liver Inflammation and Fibrosis by Regulating Tissue Cellular Immunity. Front Immunol. 2020;11:584048. doi:10.3389/fimmu.2020.584048

52. Wang H, Tsung A, Mishra L, Huang H. Regulatory T cell: a double-edged sword from metabolic-dysfunction-associated steatohepatitis to hepatocellular carcinoma. EBioMedicine. Mar 2024;101:105031. doi:10.1016/j.ebiom.2024.105031

53. Yuksel M, Wang Y, Tai N, et al. A novel "humanized mouse" model for autoimmune hepatitis and the association of gut microbiota with liver inflammation. Hepatology. Nov 2015;62(5):1536–50. doi:10.1002/hep.27998

54. Li L, Tu J, Jiang Y, Zhou J, Schust DJ. Regulatory T cells decrease invariant natural killer T cell-mediated pregnancy loss in mice. Mucosal Immunol. May 2017;10(3):613–623. doi:10.1038/mi.2016.84

55. Wang Y, Liu T, Tang W, et al. Hepatocellular Carcinoma Cells Induce Regulatory T Cells and Lead to Poor Prognosis via Production of Transforming Growth Factor-β1. Cell Physiol Biochem. 2016;38(1):306–18. doi:10.1159/000438631

56. Langhans B, Nischalke HD, Krämer B, et al. Role of regulatory T cells and checkpoint inhibition in hepatocellular carcinoma. Cancer Immunol Immunother. Dec 2019;68(12):2055–2066. doi:10.1007/s00262-019-02427-4

57. Gao Y, Guo J, Bao X, et al. Adoptive Transfer of Autologous Invariant Natural Killer T Cells as Immunotherapy for Advanced Hepatocellular Carcinoma: A Phase I Clinical Trial. Oncologist. 11 2021;26(11):e1919–e1930. doi:10.1002/onco.13899

58. Kim D, Paggi JM, Park C, Bennett C, Salzberg SL. Graph-based genome alignment and genotyping with HISAT2 and HISAT-genotype. Nat Biotechnol. Aug 2019;37(8):907–915. doi:10.1038/s41587-019-0201-4

59. Langmead B, Salzberg SL. Fast gapped-read alignment with Bowtie 2. Nat Methods. Mar 04 2012;9(4):357–9. doi:10.1038/nmeth.1923

60. Fanidis D, Moulos P. Integrative, normalization-insusceptible statistical analysis of RNA-Seq data, with improved differential expression and unbiased downstream functional analysis. Brief Bioinform. May 20 2021;22(3)doi:10.1093/bib/bbaa156

61. Gioulbasani M, Galaras A, Grammenoudi S, et al. The transcription factor BCL-6 controls early development of innate-like T cells. Nat Immunol. 09 2020;21(9):1058–1069. doi:10.1038/s41590-020-0737-y

62. Moulos P, Hatzis P. Systematic integration of RNA-Seq statistical algorithms for accurate detection of differential gene expression patterns. Nucleic Acids Res. Feb 27 2015;43(4):e25. doi:10.1093/nar/gku1273

63. Love MI, Huber W, Anders S. Moderated estimation of fold change and dispersion for RNA-seq data with DESeq2. Genome Biol. 2014;15(12):550. doi:10.1186/s13059-014-0550-8

64. Robinson MD, McCarthy DJ, Smyth GK. edgeR: a Bioconductor package for differential expression analysis of digital gene expression data. Bioinformatics. Jan 01 2010;26(1):139–40. doi:10.1093/bioinformatics/btp616

65. Ritchie ME, Phipson B, Wu D, et al. limma powers differential expression analyses for RNA-sequencing and microarray studies. Nucleic Acids Res. Apr 20 2015;43(7):e47. doi:10.1093/nar/gkv007

66. Yanming Di DWS, Jason S Cumbie and Jeff H Chang. The NBP Negative Binomial Model for Assessing Differential Gene Expression from RNA-Seq. Statistical Applications in Genetics and Molecular Biology 2011.

67. Tarazona S, Furió-Tarí P, Turrà D, et al. Data quality aware analysis of differential expression in RNA-seq with NOISeq R/Bioc package. Nucleic Acids Res. Dec 02 2015;43(21):e140. doi:10.1093/nar/gkv711

