## Supplementary figures for "Invariant Natural Killer T cells control positively and negatively the development of hepatocellular carcinoma"

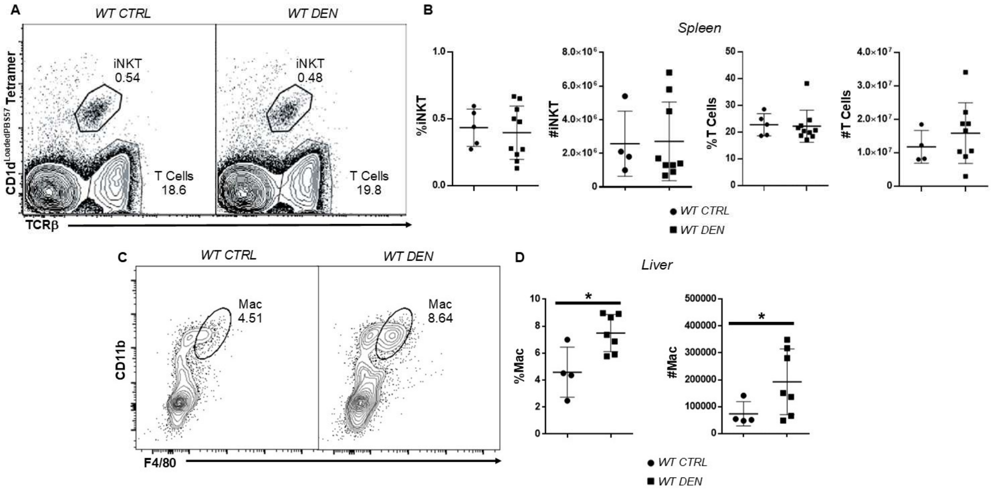


***Supplementary Figure 1. A*** *FACS plots show the frequency of iNKT and T cells in the spleen from WT CTRL and WT DEN mice.* ***B*** *Scatter plots show the percentage and numbers of iNKT and T cells from WT CTRL and WT DEN mice.* ***C*** *FACS plot show the percentage of macrophages in the liver from WT CTRL and WT DEN mice 38-42 weeks after DEN injection.* ***D*** *Scatter plots show the frequency and numbers of macrophages from WT CTRL and WT DEN mice. Bars represent mean ± SD from 5 independent experiments, with one WT and one or two WT DEN mice each. *, $p<0.05$ by t test.*

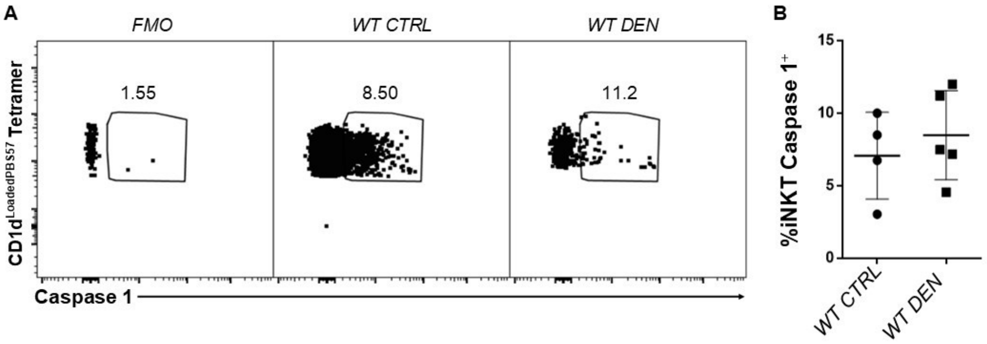


***Supplementary Figure 2. iNKT cells did not undergo pyroptosis in HCC. A*** *FACS plots show the expression of active caspase 1 on iNKT cells from WT CTRL and WT DEN mice 38-42 weeks after DEN injection.* ***B*** *Scatter plot shows the expression of active caspase 1 on iNKT cells in WT CTRL and WT DEN mice. Bars represent mean ± SD from 4 independent experiments, with one WT and one or two WT DEN mice each.*

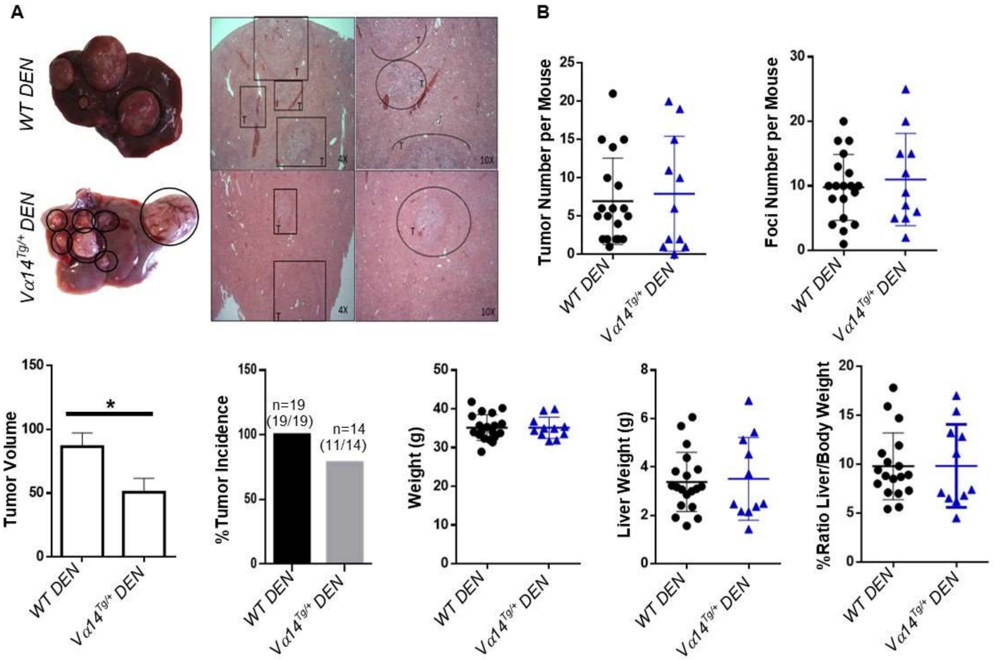


***Supplementary Figure 3. Overproduction of iNKT cells did not affect the development of HCC. A*** *Liver pictures from WT DEN (top) and V$\alpha$14*$^{Tg/+}$ *DEN (bottom) mice. H&E histology after DEN in 4X and 10X magnification,* ***T****, tumor.* ***B*** *Scatter plots show foci and tumor number, tumor size, body and liver weight (g) and %Ratio liver/body weight and bar plot shows the tumor incidence in WT DEN and V$\alpha$14*$^{Tg/+}$ *DEN mice. Bars represent mean ± SD from 11 independent experiments, with one or two WT and one or two V$\alpha$14*$^{Tg/+}$ *mice each. *, p<0.05 by t test.*

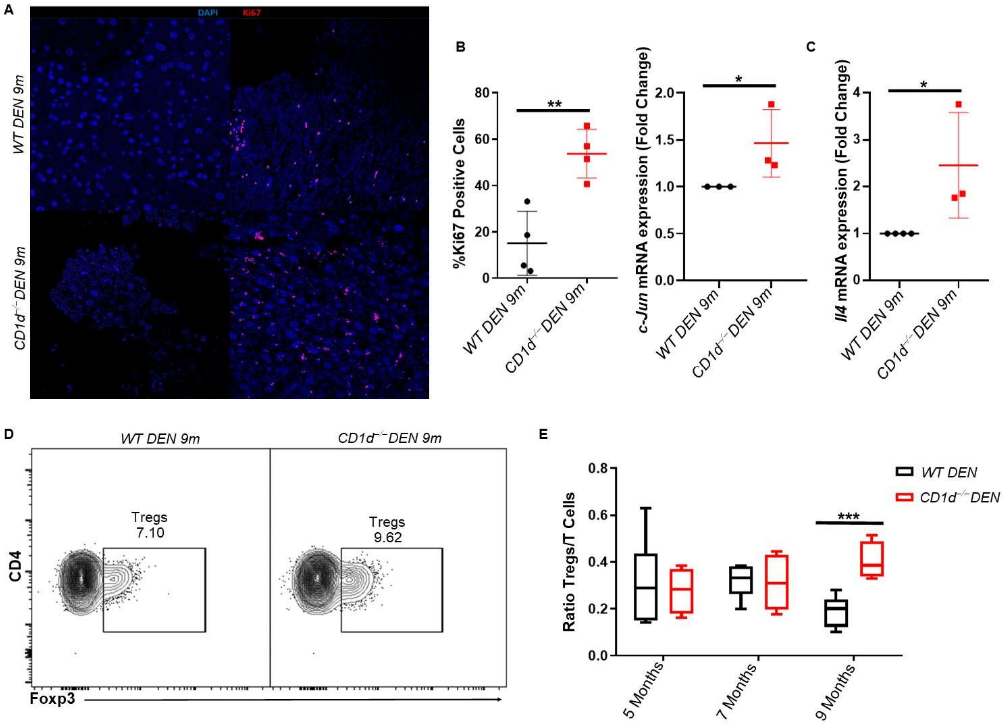


***Supplementary Figure 4. A*** *Representative immunofluorescence images of WT DEN (top) and CD1d$^{-/-}$ DEN (bottom) livers stained with anti-Ki67 (red) and co-labeled with DAPI (blue) (right). Negative control of anti-Ki67 is shown (left).* ***B*** *Scatter plot shows the percentage of Ki67 positive cells. Scatter plot shows the mRNA expression of the proliferation marker, c-Jun, in WT DEN and CD1d$^{-/-}$ DEN.* ***C*** *Scatter plot shows the mRNA expression levels of anti-inflammatory cytokine, Il4, in WT DEN and CD1d$^{-/-}$ DEN.* ***D*** *FACS plots show the frequency of Tregs in WT DEN and CD1d$^{-/-}$ DEN.* ***E*** *Scatter plot show the Ratio of Tregs/T cells in WT DEN and CD1d$^{-/-}$ DEN 9 months after DEN injection. Bars represent mean ± SD from 3 independent experiments, with one or two WT DEN and CD1d$^{-/-}$ DEN mice in each time point. *, $p<0.05$, **, $p<0.01$ and ***, $p<0.001$ by t test.*

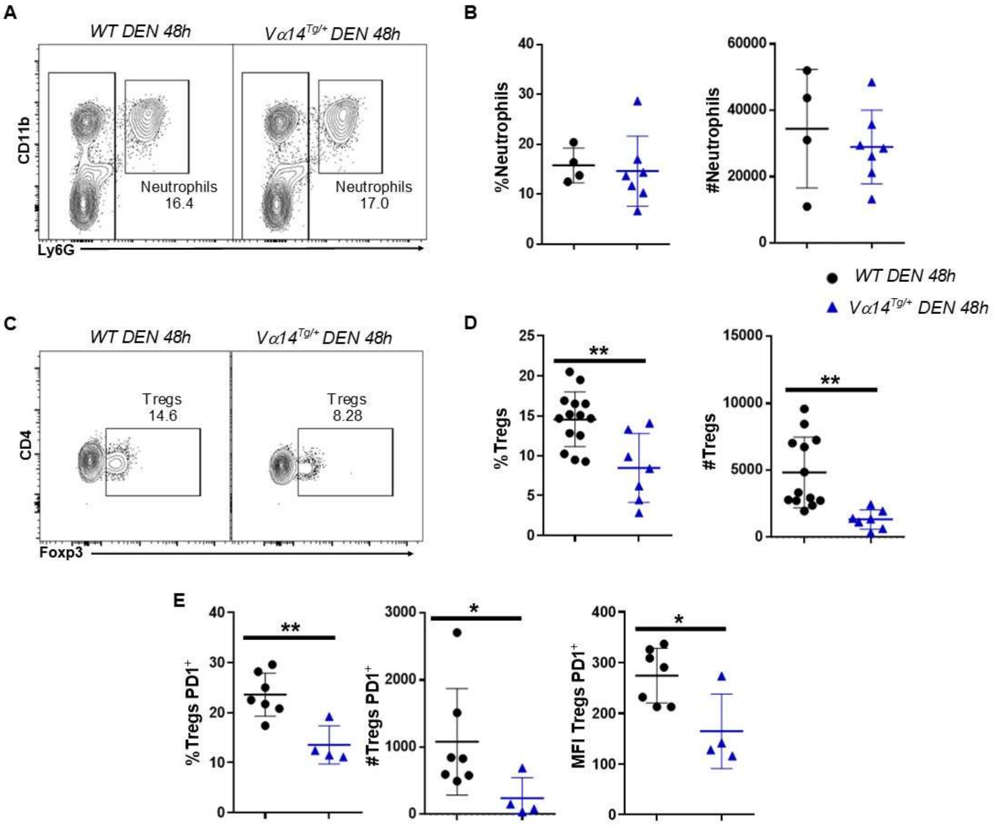


***Supplementary Figure 5.*** ***A*** *FACS plots show the frequency of neutrophils ($CD11b^+Ly6G^+$) in WT DEN and $V\alpha14^{Tg/+}$ DEN mice 48 hours after DEN injection.* ***B*** *Scatter plots show the percentage and number of neutrophils in WT DEN and $V\alpha14^{Tg/+}$ DEN mice.* ***C*** *FACS plots show the $TCR\beta^+CD4^+Foxp3^+$ lymphocyte population from WT DEN and $V\alpha14^{Tg/+}$ DEN.* ***D*** *Scatter plots show the frequency and number of Tregs.* ***E*** *Scatter plots show the frequency and number of Tregs positive to PD1 and the MFI of the expression of PD1 on Tregs in WT DEN and $V\alpha14^{Tg/+}$ DEN 48 hours post-DEN administration. Bars represent mean ± SD from at least 3 independent experiments, with one or two WT DEN and one or two $V\alpha14^{Tg/+}$ DEN mice each. *, $p<0.05$, **, $p<0.001$ by t test.*

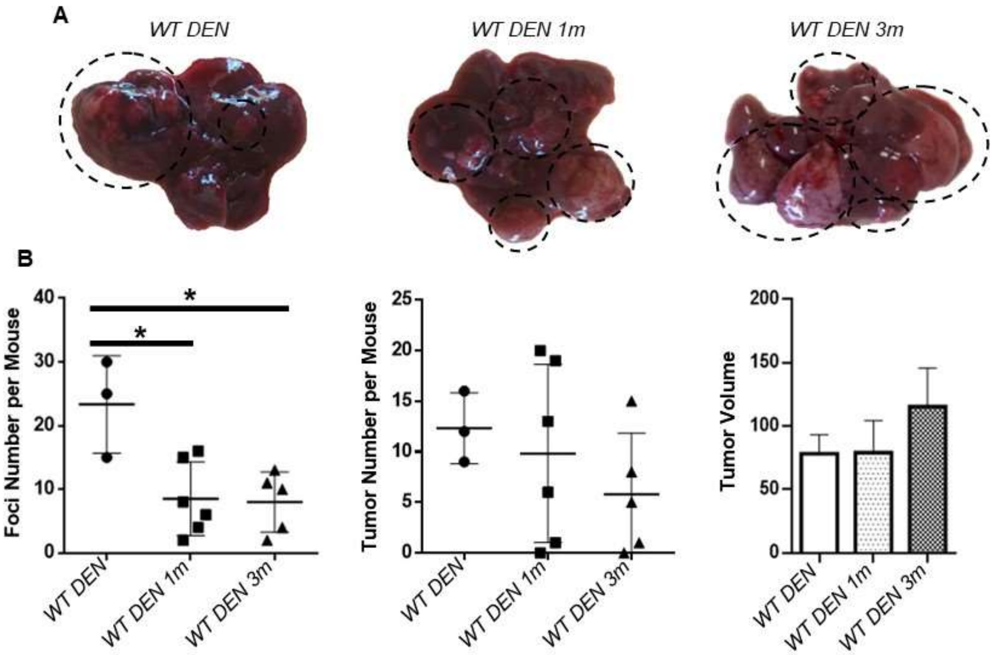


***Supplementary Figure 6.*** ***A*** *Liver images from WT DEN without iNKT cells adoptive transfer (right), WT DEN with iNKT cells adoptive transfer 1 month after DEN injection (middle) and WT DEN with iNKT cells adoptive transfer 3 months post-DEN injection (left).* ***B*** *Scatter plots show the number of foci and tumors per mouse and bar plot shows the tumor volume among WT DEN without iNKT cells adoptive transfer, and WT DEN with adoptive transfer of iNKT cells in 1– and 3– months post-DEN administration. Bars represent mean ± SD from 3 independent experiments, with one WT DEN and one or two WT DEN mice each. *, p<0.05 by t test.*
